# A cross-center comparison of the relationship between matriline fragmentation, grooming cohesion, and agonistic behavior in captive rhesus macaque (*Macaca mulatta*) social groups

**DOI:** 10.64898/2026.01.18.700196

**Authors:** B.A. Beisner, B. McCowan, M.A. Bloomsmith, L. Lacefield, K. F. Ethun

## Abstract

A major challenge in managing captive-bred rhesus macaque social groups is mitigating deleterious aggression before it escalates to social instability. Prior work at the California National Primate Research Center (CNPRC) showed that fragmentation of matrilineal structure—reflected in lower average kinship among female kin—is associated with weakened cohesion in grooming networks and higher rates of intense aggression. We tested the generality of these findings by analyzing data from 105 matrilines across 16 social groups at CNPRC and Emory NPRC (ENPRC), which differ in group size, natal male management, and housing. Using generalized linear models, we found that matrilines with lower mean kinship coefficients showed greater grooming fragmentation, even after accounting for network density. Threshold analyses identified a mean kinship of 0.16 as the point at which grooming cohesion declined most consistently across both centers, highlighting a biologically meaningful level of relatedness for maintaining kin-biased social bonds. Patterns of severe aggression differed by target and center: across both centers, matrilines with lower mean kinship directed proportionally more severe aggression toward kin. However, for aggression toward all group members, lower kinship predicted more severe aggression only at ENPRC; at CNPRC, this effect emerged only when natal male aggression was included. Our results demonstrate that mean matrilineal kinship is a robust indicator of family cohesion and latent social instability across management settings. Nepotistic threshold analysis provides a practical tool for managers to identify matrilines at risk for social fragmentation and implement interventions before intra-family aggression emerges.

## Introduction

Affiliative relationships are a core feature of primate societies and contribute directly to individual health, welfare, and reproductive success. In both wild and captive settings, primates use grooming and other affiliative interactions to regulate stress, support psychological well-being, and maintain social bonds (Boccia, Reite & Laudenslager, 1989; Aureli, Preston & de Waal, 1999; Silk, Alberts & Altmann, 2003; Shutt et al., 2007; Silk et al., 2010). These relationships also promote group-level cohesion (Lehmann, Korstjens & Dunbar, 2007), underscoring the importance of rich social environments in captivity and why social housing is considered the most effective form of enrichment (Lutz & Novak, 2005).

In many primates, social relationships are shaped by patterns of kinship. In Old World monkeys, maternal kin often form the core of social groups, and females show strong affiliative biases toward close relatives (Kurland, 1977; Massey, 1977; Kaplan, 1978; Seyfarth & Cheney, 1984; Lopez-Vergara & Santillan-Doherty, 1989; Silk, 2001). These kin-based relationships influence proximity, grooming, conflict intervention, and alliance formation, which reinforce dominance hierarchies and promote social stability (Reviewed by: Gouzoules 1984; Gouzoules & Gouzoules 1987; Walters 1987; Kapsalis & Berman 1996; Silk 2001). In rhesus macaques, groups are organized around steep, nepotistic dominance hierarchies, with strong maternal kin bias. Mothers and other close kin actively support the rank acquisition of younger females, such that daughters inherit a rank just below their mother (Missakian, 1972; Sade, 1972; Berman, 1980; Chapais, 1992).

Kin bias, however, is not distributed equally across all relatives. Affiliative and supportive behaviors (e.g., grooming, agonistic intervention) are most frequent among close kin, but these biases diminish beyond what has been described as a “nepotistic threshold”. Importantly, this threshold is not a single fixed value but instead falls within the range of r = 0.25–0.125, at which point the rates of interactions among distant kin resemble those between nonkin (Massey, 1977; Kaplan, 1978; Kapsalis & Berman, 1996a; Rendall et al., 1996; Chapais et al., 1997, 2001). Moreover, the precise point appears to vary by relationship type: in Japanese macaques, for example, Chapais et al. (2001) found that adult females were more likely to intervene in conflicts involving direct kin (e.g., mothers, grandmothers, great-grandmothers) at relatedness levels as low as r = 0.125, whereas support for collateral kin (e.g., sisters, aunts) had an upper limit of r = 0.25.

When family structures are disrupted, social instability can arise, including outbreaks of deleterious aggression and social overthrows (Gygax, Harley & Kummer, 1997; Oates-O’Brien et al., 2010; McCowan, Beisner & Hannibal, 2017). Earlier work has demonstrated that matrilines with lower mean kinship coefficients, indicative of greater pedigree fragmentation, exhibited less cohesion in grooming networks, higher rates of within-family aggression, and increased risk of trauma (Beisner et al., 2011a). Subsequent studies have shown similar connections between matriline structure and social stability. For example, Johnston et al. (2020) found that an outbreak of intrafamily aggression, which involved distant kin from separate matriline branches (mean kinship < 0.20), was preceded by a decline in food intake. Stavisky et al. (Stavisky et al., 2018) found that more recently formed groups, composed of many small matrilines, had higher trauma rates compared to longstanding groups with larger, multigenerational matrilines, again suggesting a link between matriline structure and group stability.

Social stability in rhesus groups can be shaped not only by internal dynamics but also by institutional management practices—especially those that influence group composition. One key area of variation is how natal males are managed, a factor that affects both kin structure and social cohesion. Some facilities retain some natal males in their birth groups into adulthood (Smith, 1986; Beisner et al., 2011b; Pritchard et al., 2024), while others routinely remove them around puberty. In some cases, new adult males are introduced periodically to replace natal males and maintain genetic diversity for breeding (Rox et al., 2019, 2021; Bailey et al., 2021b,a). Retaining natal males preserves close kin ties with mothers and sisters (Beisner et al., 2011b), whereas removal might redirect maternal social investment toward more distantly related females, potentially buffering against matrilineal fragmentation.

Alongside natal male management practices, institutional variation in available housing, such as differences in enclosure design or capacity, can influence group size and, in turn, shape the structure and expression of affiliative relationships. In both wild and captive primates, group size can impose practical limits on grooming behavior. Even when cognitive capacity is not exceeded, time constraints arising from ecological demands such as foraging or vigilance may reduce the time available to maintain social bonds (Dunbar, 1992; Dunbar 2004; Lehmann, Korstjens & Dunbar, 2007; Dunbar, Korstjens, Lehmann 2009). When groups become too large, individuals may not have sufficient time to groom all potential partners, and cohesion can decline. When ecological constraints are relaxed in captivity, individuals in small groups may be able to distribute grooming broadly across kin, creating apparent cohesion even within fragmented matrilines.

Although higher average matrilineal kinship is associated with stronger cohesion in rhesus groups (Beisner et al., 2011a), it is unclear whether this association generalizes across colonies with different structures and management practices. Here, we extend earlier work by testing whether the kinship–cohesion relationship still holds in breeding groups at ENPRC, which differ from CNPRC in three ways: smaller average group size, fewer matrilines per group, and systematic natal male removal. We also introduce a novel approach to identifying a matriline-level nepotistic threshold, using a series of kinship cutoff models to estimate the point at which grooming cohesion declines. Specifically, we hypothesized that matriline cohesion would remain relatively stable until average kinship fell below a critical threshold, after which cohesion would weaken sharply. We further hypothesized that matrilines with lower mean kinship would show higher rates of severe aggression, and that the strength of these relationships would vary across centers. By comparing two NPRCs with distinct social demographics and management policies, we aim to clarify the generality of these patterns and provide updated guidance for breeding colony management to minimize kinship fragmentation and social unrest.

## METHODS

### Ethics Statement

All research procedures adhered to the American Society of Primatologists Principles for the Ethical Treatment of Nonhuman Primates (ASP, 2021), the ASP Code of Best Practices for Field Primatology (ASP, 2014, where relevant), the US National Research Council’s *Guide for the Care and Use of Laboratory Animals* (2011), the US Public Health Service’s Policy on Humane Care and Use of Laboratory Animals, and all applicable US legal requirements.

### Artificial Intelligence Disclosure

The first author used the AI tool ChatGPT (OpenAI) as an aid during the writing process. ChatGPT assisted with reviewing and revising manuscript text, suggesting alternative phrasing, clarifying wording, and providing feedback on sentence and paragraph structure. The tool was not used for data collection, analysis, figure generation, or literature search and interpretation. All content and interpretations remain the sole responsibility of the authors.

Research at the California National Primate Research Center (CNPRC; University of California, Davis) was approved by the UC Davis Institutional Animal Care and Use Committee. All methods were observational, involved no experimental or invasive procedures, and subjects were housed in large outdoor social groups in accordance with CNPRC colony management practices. CNPRC is fully accredited by AAALAC International.

Research at the Emory National Primate Research Center (ENPRC; formerly Yerkes NPRC) was approved by the Emory University Institutional Animal Care and Use Committee. All ENPRC studies were either purely observational or coordinated with existing colony management practices (e.g., breeder male introductions), and no procedures beyond standard husbandry and care were performed. ENPRC is also fully accredited by AAALAC International.

### Study Site and Groups

The subjects were group-living rhesus macaques (*Macaca mulatta*) housed at two National Primate Research Centers (CNPRC and ENPRC). Behavioral data came from four studies: one at CNPRC and three at ENPRC.

#### CNPRC

We studied seven multimale–multifemale groups in large 1,800 m² outdoor enclosures between June 2008 and December 2009. Groups ranged from 105–198 animals and comprised 6–14 matrilines each (Table 1). Enclosures were equipped with multiple A-frame houses, perches, swinging barrels, and other enrichment. All groups had continuous access to drinking water and were fed a commercial monkey diet twice daily, supplemented with fresh produce and scratch. Matrilines with at least four adult females (≥3 years old) were included, yielding 47 CNPRC matrilines.

**Table 1.** Study group characteristics and observation hours.

| Center | Study | Group ID | Adults (3+ yrs) | Group Size | Matriline N $\geq 4$ F | Total Obs Hours |
| --- | --- | --- | --- | --- | --- | --- |
| CNPRC | #1 - June 2008 - October 2009 | 1 | 99 | 176 | 11 | 162 |
|  |  | 10 | 85 | 164 | 5 | 174 |
|  |  | 14 | 54 | 108 | 6 | 144 |
|  |  | 16 | 77 | 150 | 6 | 144 |
|  |  | 18 | 99 | 198 | 6* | 144 |
|  |  | 5 | 82 | 137 | 8* | 168 |
|  |  | 8 | 91 | 160 | 7* | 144 |
| ENPRC | #2 March | I | 42 | 84 | 4 | 163 |
|  | 2017 -<br>December<br>2019 | II | 26 | 64 | 6 | 152 |
|  |  | II | 33 | 76 | 4 | 168 |
|  |  | V | 16 | 41 | 2* | 159 |
|  |  | VI | 18 | 44 | 1* | 163 |
|  |  | VII | 34 | 60 | 2* | 168 |
|  | #3<br>September<br>2021 - July<br>2022 | II | 37 | 81 | 4 | 115 |
|  |  | IX | 45 | 94 | 3 | 112 |
|  |  | VIII | 80 | 181 | 11 | 150 |
|  |  | X | 26 | 67 | 1* | 83 |
|  | #4<br>September<br>2023 - May<br>2025 | I | 60 | 106 | 6 | 480 |
|  |  | VIII | 79 | 153 | 8* | 528 |
|  |  | XI | 64 | 121 | 4* | 528 |

\*Group has multiple additional matriline, but they were too small for analysis (i.e., <4 adult members)

Behavioral data were collected by two observers using standardized methods. Agonistic interactions (e.g., displacements, threats, lunges, chases, bites) were recorded with event sampling, noting initiator, recipient, and behavioral sequence. Affiliative interactions (grooming and social contact) were recorded via scan sampling every 20 minutes. Each group was observed six hours per day, four days per week, on a four-week rotating schedule, yielding at least 144 observation hours per group. These data were previously published (Beisner et al. 2011a) and are reanalyzed here alongside ENPRC data.

#### ENPRC

At the Field Station (Lawrenceville, GA), data were collected from nine unique social groups across three studies conducted between March 2017 and May 2025 (Study #2: 2017–2019; Study #3: 2021–2022; Study #4: 2023–2025). Some groups were studied more than once (Groups I, II, VIII) with 2–7 years between samplings. Groups ranged from 33–181 animals (17–80 adults), encompassing 3–11 matrilines per group (Table 1). ENPRC group sizes were generally smaller than CNPRC groups (Figure 1a).

**Figure 1.**
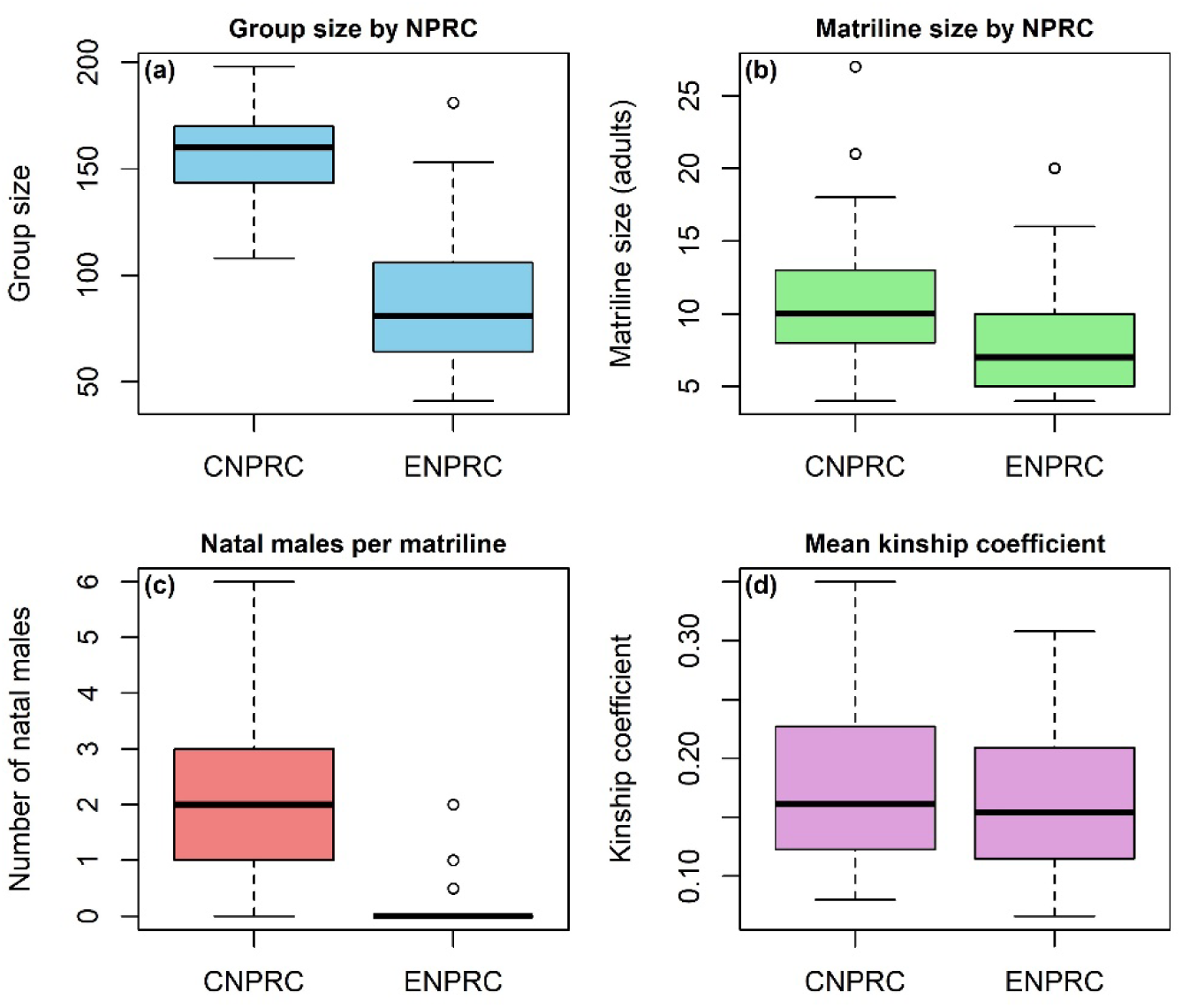
Comparison of group- and matriline-level demographics across the two NPRCs: group size (a), matriline size (b), number of natal males per matriline (c), and mean matrilineal kinship coefficient (d)

All groups lived in indoor–outdoor compounds with outdoor areas of 232–1,450 m² and indoor areas of 15–35 m². Animals had continuous access to drinking water and were either fed twice daily or given unrestricted access to a commercial diet via automated feeders (Johnston et al. 2020). Enrichment included fresh produce, climbing structures, foraging devices, and manipulanda. All ENPRC animals were SPF (free of SIV, simian T-lymphotropic virus, simian type D retroviruses, and herpes simian B virus). Matrilines with at least four adult females were included, yielding 58 ENPRC matrilines.

Behavioral data collection protocols were designed or co-designed by the first author to match CNPRC methods. For all studies, 1–2 observers recorded agonistic and affiliative interactions among animals aged ≥3 years. Agonistic behavior was recorded using event sampling, and affiliation with scan sampling every 20–30 minutes. Most ENPRC groups were smaller than CNPRC groups (Table 1), allowing a single observer to collect both event- and scan-sampling data concurrently.

In Study #2, groups were observed 3–6 hours/day for 31–37 days across six months (150–168 hours per group). In Study #3, groups were observed 2–4 hours/day for 22–40 days across four months (85–150 hours per group). In Study #4, groups were observed 12 hours/week for 10 months, yielding 480–528 hours per group. Inter-observer reliability tests confirmed ≥95% accuracy for individual ID and ≥85% reliability for behavior coding. Data were recorded using the HandBase app on handheld devices, or by voice recorder, if needed, and then transcribed.

### Matrilines

A matriline was defined as all living group members descended from a single female common ancestor (the matriarch) who was a founding group member. Study groups ranged from 6–32 years old (CNPRC: 6–32 years, mean = 20.3; ENPRC: 7–26 years, mean = 20.0) and contained matrilines spanning 3–7 generations. Matriline sizes were comparable between centers (Figure 1b), but group sizes and natal male numbers differed more markedly (Figure 1c).

For each matriline, we calculated the average kinship coefficient among all pairs of adult females, following Beisner et al. (2011a). For example, mother–daughter pairs have a coefficient of 0.5, grandmother–granddaughter or half-siblings 0.25, and so on. Higher mean kinship indicates proportionally more close-kin dyads. In our data, mean kinship ranged from 0.067–0.30 (mean = 0.17, SD = 0.05) for ENPRC and 0.08–0.35 (mean = 0.18, SD = 0.07) for CNPRC (Figure 1d). Only matrilineal kinship was considered; paternal ties were excluded.

### Grooming Networks

We constructed directed, weighted grooming networks for each study group using all recorded interactions among adult females (≥3 years). Male interactions were excluded to enable comparisons between centers with different male-management practices. At ENPRC, most males are removed by age three, whereas at CNPRC many remain in their natal groups into adulthood (Figure 1c). To test for male effects, we included the number of natal males per matriline as a predictor in our models.

Network density—the proportion of possible ties observed— was calculated using the *igraph* package in R (Csardi & Nepusz, 2006). Grooming rates and densities were slightly higher at ENPRC (0.07–0.14 interactions/hour; density 0.20–0.63) than CNPRC (0.05–0.08; density 0.11– 0.20). Prior work demonstrated that CNPRC networks are dense enough for reliable community detection (Beisner et al., 2011a), indicating that ENPRC networks also meet this criterion.

Because higher density can reduce modularity and obscure community structure, we addressed this in two ways: (1) including edge density as a covariate, and (2) constructing density-matched ENPRC networks by subsampling consecutive observation sessions until edge density was ≤0.20 while retaining all individuals. For two groups (V and X), a density of 0.20 could not be reached without excluding nodes; in these cases, we used the lowest possible density that preserved the full group (Table 2).

**Table 2.** Grooming network metrics for each study group (networks included only female-female grooming interactions)

| Study | Group ID | Total Grooming | Edge Density | Groom Communities | Network Modularity |
| --- | --- | --- | --- | --- | --- |
| #1<br>CNPRC | 1 | 570 | 0.15 | 10 | 0.51 |
|  | 10 | 912 | 0.19 | 11 | 0.39 |
|  | 14 | 285 | 0.20 | 8 | 0.41 |
|  | 16 | 361 | 0.15 | 11 | 0.33 |
|  | 18 | 560 | 0.11 | 8 | 0.53 |
|  | 5 | 389 | 0.13 | 9 | 0.38 |
|  | 8 | 556 | 0.12 | 14 | 0.47 |
| #2<br>ENPRC | I | 668 | 0.32 | 4 | 0.49 |
|  | II | 538 | 0.43 | 6 | 0.44 |
|  | II | 600 | 0.35 | 5 | 0.48 |
|  | V | 353 | 0.63 | 4 | 0.31 |
|  | VI | 224 | 0.46 | 3 | 0.28 |
|  | VII | 427 | 0.37 | 6 | 0.48 |
| #3<br>ENPRC | II | 407 | 0.27 | 5 | 0.44 |
|  | IX | 670 | 0.26 | 8 | 0.53 |
|  | VIII | 1322 | 0.20 | 12 | 0.43 |
|  | X | 262 | 0.24 | 5 | 0.46 |
| #4<br>ENPRC | I | 2856 | 0.43 | 5 | 0.41 |
|  | VIII | 3960 | 0.42 | 11 | 0.30 |
|  | XI | 2984 | 0.37 | 8 | 0.43 |

To visualize the density–modularity relationship, we plotted modularity against density for each network with fitted lines by center (Supplementary Figure S1). This confirmed a strong negative relationship and justified both the statistical control and density-matched analyses. Subsampling results for ENPRC groups are provided in Supplementary Figures S2–S20.

### Community Structure

We identified grooming communities using the Walktrap algorithm in *igraph* (Pons & Latapy, 2005; Csardi & Nepusz, 2006), which iteratively merges nodes into clusters based on short random walks. This method incorporates weighted edges and has been shown to reliably detect community structure across a range of network sizes and mixing parameters (Yang, Algesheimer & Tessone, 2016; Smith et al., 2020). The optimal partition is the one that maximizes modularity, which quantifies the extent to which edges are denser within clusters than between them (Newman & Girvan, 2004).

Because rhesus macaque females preferentially groom close kin (Sade, 1972; Berman & Kapsalis, 1999; Silk, 2006), we expected matriline members to cluster together within grooming communities. To quantify matriline fragmentation, we used the group-level grooming network to identify each matriline’s *primary grooming community* (i.e., the one containing the largest proportion of its members) and then counted the number of females who fell outside that cluster. This measure captures cohesion relative to the *whole-group* structure, rather than within isolated matrilineal subnetworks. Our previous approach (Beisner et al., 2011a) constructed separate networks for each matriline and excluded small lineages (<5 adult females) to ensure sufficient size. The present approach, by contrast, defines fragmentation relative to group-level community structure, which allows the inclusion of smaller matrilines. To balance inclusivity with analytical robustness, we restricted analyses to matrilines with ≥4 adult females, and we included matriline size as an offset in all statistical models.

### Handling of Animal Removals

Several animals were permanently removed during the study for research or clinical reasons: 22 subadult natal males (CNPRC: N=19, ENPRC: N=3) and 10 adult females (CNPRC: N=3, ENPRC: N=7). To ensure that such removals did not bias the grooming cohesion analysis, females present for <66% of the observation period were excluded from the outcome variable but retained in grooming networks to represent existing social ties accurately.

#### Aggressive Behavior and Matriline Rank Variables

To investigate whether more fragmented matrilines also experienced greater internal conflict or higher overall participation in severe aggression, we calculated two measures of the use of severe aggression. First, we determined the proportion of all aggression initiated by the matriline that was severe (i.e., involved biting or pinning the opponent to the ground). Second, we calculated the proportion of aggression directed toward kin that was severe.

All agonistic interactions with clearly defined winners and losers (e.g., aggression met with only submissive behavior; unprovoked approach resulting in submission) were analyzed using the *Perc* package (Fujii et al., 2014) in RStudio to generate a dominance hierarchy among adult females in each study group. Because females from the same matriline typically occupy adjacent ranks (Sade, 1969), matrilines were assigned a rank order based on the relative positions of their members. To standardize matriline rank for statistical analyses, we calculated the proportion of other matrilines outranked by each matriline (range: 0 [lowest rank] to 1 [highest rank]). Matrilines with no recorded aggression involving kin were excluded from the within-family severe aggression model (final n = 103 matrilines), as the proportion could not be defined.

#### Statistical Analysis

We analyzed data from 105 matrilines using generalized linear models (GLMs) to test the relationship between matriline kinship structure, kin-based grooming cohesion, and aggression severity across two research institutions with different group management practices. We quantified kin-based grooming cohesion as the number of adult females (≥3 years old) in each matriline who clustered outside the matriline’s primary grooming community within the whole-group network. Because this count is constrained by matriline size, we included an offset term for the total number of adult females per matriline. We ran two sets of models: one using the full networks with network density included as a covariate, and another using density-matched networks. Preliminary checks showed mild overdispersion for some models, so we compared Poisson and negative binomial families and used the negative binomial when it improved model fit and residual dispersion.

Aggression severity was assessed using two related outcomes: (1) the proportion of all aggression initiated by each matriline that was severe (regardless of target), and (2) the proportion of kin-directed aggression that was severe. For both outcomes, we modeled the number of severe versus non-severe conflicts using a quasibinomial family to handle moderate overdispersion (Bolker, 2015). Predictors included mean kinship coefficient, matriline rank (standardized as the proportion of other matrilines outranked), group size, number of natal males per matriline, and NPRC, along with the interaction term kinship × NPRC to test whether effects differed between centers.

When possible, we used AICc (Akaike’s Information Criterion corrected for small sample size) to select the most parsimonious models, retaining all models with ΔAICc < 2 (Burnham & Anderson, 2002; Burnham, Anderson & Huyvaert, 2011). For quasibinomial models, AIC is not available, so we compared nested models using deviance-based F-tests to determine whether adding predictors significantly improved model fit, following standard practice in R’s **anova.glm()** and **drop1(…, test=“F”)** routines (Hastie & Pregibon, 1992). Analyses were conducted in R version 3.6.1 (R Core Team, 2019) using the *MASS* and *stats* packages for Poisson and negative binomial models. We also tested random effects of social group identity using the *lme4* package (Bates et al., 2013), but these did not improve model fit and were omitted from final models.

To further explore whether a kinship-based threshold of social cohesion exists at the matriline level, we constructed binary versions of the mean kinship coefficient, coding matrilines as *highly related* (=1) when their mean coefficient was greater than or equal to a given cutoff and *distantly related* (=0) otherwise. We tested thresholds ranging from 0.25 to 0.10, spanning the range of nepotistic thresholds reported in the primate literature. For grooming cohesion, we substituted each binary kinship term into the best-fitting GLMs and compared AICc values across models to identify which threshold best explained the data. To parallel this approach, we also tested whether binary kinship thresholds better explained aggression outcomes. Because these models required quasibinomial error structures, we evaluated model fit using deviance-based F-tests rather than AICc, comparing each threshold model against the corresponding continuous kinship model.

## RESULTS

The grooming networks for all study groups showed moderate to strong levels of modularity, ranging from 0.28 to 0.53 (Table 2), indicating clear community structure. Matriline members tended to cluster together in the same community. On average, 2.23 females per matriline (±2.63 SD; range: 0–14) clustered separately from their matriline, representing 24.1% of matriline members on average (range: 0–75%). Grooming communities contained an average of 1.63 matrilines (±1.27 SD; range: 1–7 matrilines).

### Mean kinship coefficient predicts matriline social cohesion

Analyses of the full dataset showed that matrilines with lower mean kinship coefficients had more females clustering away from their family’s main grooming community (β = –5.67, SE = 1.58, *p* < 0.001; Figure 2). Larger group size (β = –0.011, SE = 0.005, *p* = 0.027), lower edge density (β = –2.75, SE = 0.97, *p* = 0.005), and lower network modularity (β = –3.26, SE = 1.27, *p* = 0.010) were also associated with greater matrilineal grooming cohesion. Including the interaction NPRC × mean kinship did not improve model fit (Table 3), indicating that the relationship between kinship and cohesion was consistent across centers.

**Figure 2.**
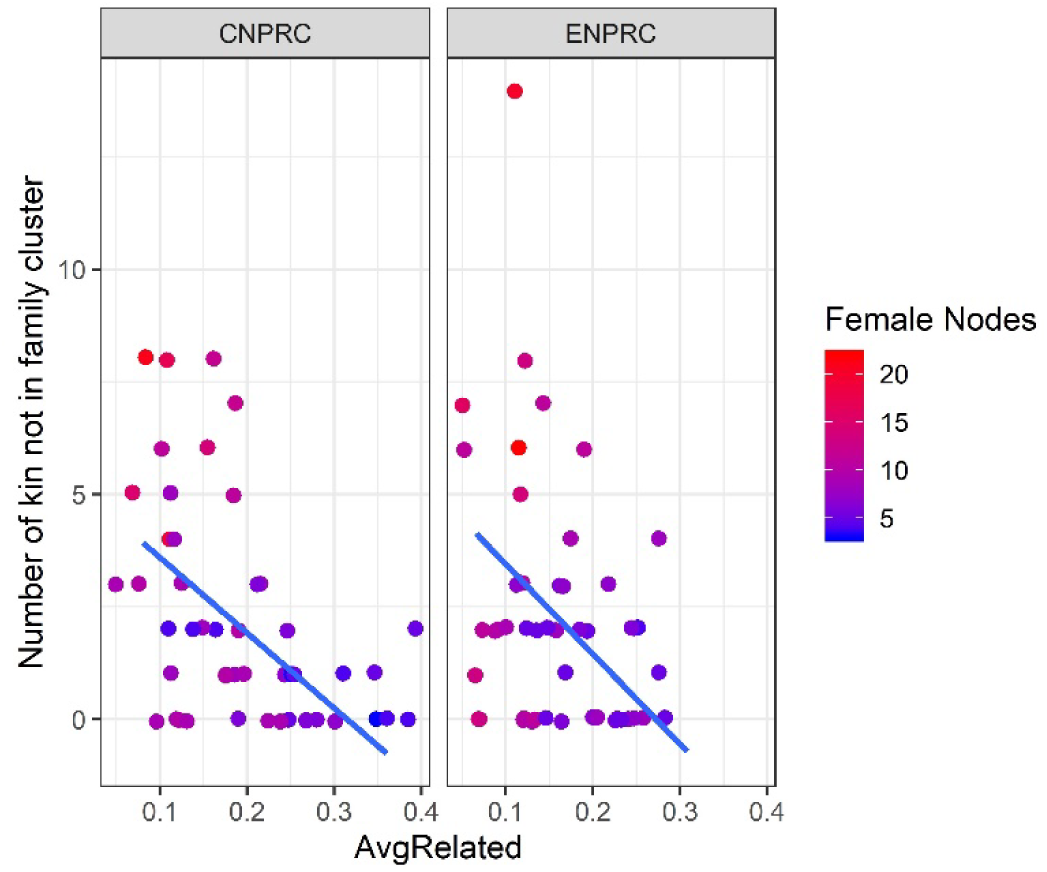
The number of females that clustered separately from their matriline plotted by mean kinship coefficient.

**Table 3.** Top four candidate models predicting the number of adult females per matriline clustering outside their primary grooming community, with model equations and AICc values, for full networks.

| Model Number | Model Equation | AICc | $\Delta$ AIC |
| --- | --- | --- | --- |
| 1 | NumKinClusterAway ~ MeanKinship + Density + Modularity + Group Size | 367.8 | 0 |
| 2 | NumKinClusterAway ~ MeanKinship + NPRC + Density + Modularity + Group Size | 370.0 | 2.2 |
| 3 | NumKinClusterAway ~ MeanKinship*NPRC + Density + Modularity + Group Size | 372.2 | 4.3 |
| 4 | NumKinClusterAway ~ Density + Modularity + Group Size | 379.1 | 11.3 |

Analyses using the density-matched dataset — in which ENPRC networks were subsampled to match CNPRC edge densities — yielded similar results. The best-fit model included mean kinship coefficient and matriline rank. Again, higher mean kinship coefficient predicted greater cohesion (β = –4.67, SE = 1.38, *p* < 0.001). Higher matriline rank was also associated with greater cohesion (β = –0.49, SE = 0.25, *p* = 0.048). Including the interaction NPRC × mean kinship did not substantially improve model fit (Table 4).

**Table 4.** Top for candidate models predicting the number of adult females per matriline clustering outside their primary grooming community, with model equations and AICc values, for density-matched grooming networks.

| Model Number | Model Equation | AICc | $\Delta$ AIC |
| --- | --- | --- | --- |
| 1 | NumKinClusterAway ~ MeanKinship + MatrilineRank | 364.5 | 0 |
| 2 | NumKinClusterAway ~ MeanKinship*NPRC + GroupSize | 364.8 | 0.4 |
| 3 | NumKinClusterAway ~ MeanKinship*NPRC +<br>MatrilineRank | 365.0 | 0.5 |
| 4 | NumKinClusterAway ~ MeanKinship + NPRC +<br>MatrilineRank | 366.6 | 2.1 |

#### Severe Aggression and Kinship

Unlike the consistent effect of kinship on grooming cohesion, the association between mean kinship coefficient and severe aggression varied by center and by the presence of natal males. We examined two outcomes: (1) the proportion of all aggression initiated that was severe (regardless of target), and (2) the proportion of kin-directed aggression that was severe.

### Proportion of All Aggression That Was Severe

An analysis of deviance showed that matriline rank significantly explained variation in the proportion of aggression that was severe (F□,□□□ = 94.18, p < 0.0001). Adding mean kinship coefficient, NPRC, their interaction, group size, and number of natal males significantly improved model fit (F□,□□ = 9.15, p < 0.0001). The kinship × NPRC interaction was significant, indicating different patterns across centers. The model output is shown in Table 5.

**Table 5.** Coefficients and significance for the best-fit model predicting the proportion of all aggression initiated by a matriline that was severe.

| Predictor | Estimate | SE | t-value | p-value |
| --- | --- | --- | --- | --- |
| Intercept | -2.184 | 0.272 | -8.024 | <0.001 |
| Mean Kinship | -1.019 | 1.107 | -0.921 | 0.360 |
| NPRC (ENPRC) | 0.236 | 0.240 | 0.981 | 0.329 |
| Group Size | -0.0077 | 0.0025 | -3.110 | 0.002 ** |
| Number of Natal Males | 0.061 | 0.035 | 1.768 | 0.080 |
| Matriline Rank | 1.204 | 0.147 | 8.168 | <0.001 *** |
| Mean Kinship × NPRC | -4.018 | 1.481 | -2.713 | 0.008 ** |

**Note.** Model fit: quasibinomial GLM. Full model significantly improved fit compared to the intercept-only model ( $F_{6,98} = 9.15, p < 0.001$ ).

At ENPRC, matrilines with lower mean kinship initiated a significantly higher proportion of severe aggression (p = 0.008), whereas at CNPRC, mean kinship was not significantly associated with severe aggression (p = 0.36). Beyond kinship, higher-ranking matrilines (β = 1.20, SE = 0.15, p < 0.0001) and those in smaller groups (β = –0.008, SE = 0.002, p = 0.002) initiated proportionally more severe aggression. There was also a trend for matrilines with more natal males to do so (β = 0.061, SE = 0.035, p = 0.080).

To clarify the null CNPRC finding, we re-analyzed the original dataset (Beisner et al., 2011a) using a quasibinomial model. When including both female- and natal male–initiated aggression, lower mean kinship significantly predicted higher rates of severe aggression (p = 0.006), replicating the original result. However, when restricted to female-initiated aggression, the effect weakened and was only marginally significant (p = 0.09). These results suggest that natal males accounted for much of the earlier CNPRC effect.

### Proportion of Aggression Toward Kin That Was Severe

For the subset of aggression directed at kin, a model including matriline rank, mean kinship coefficient, and NPRC significantly improved fit over the null model (F _3,101_ = 9.96, *p* < 0.001). Adding the kinship × NPRC interaction, group size, and number of natal males provided only marginal additional explanatory power (F _3,98_ = 2.21, *p* = 0.092), so we focus on the more parsimonious model (Table 6).

**Table 6.** Coefficients and significance for the best-fit model predicting the proportion of kin-directed aggression that was severe

| Predictor | Estimate | SE | t-value | p-value |
| --- | --- | --- | --- | --- |
| Intercept | -1.912 | 0.195 | -9.821 | <0.001 *** |
| Matriline Rank | 0.524 | 0.167 | 3.135 | 0.002 ** |
| Mean Kinship | -3.214 | 0.986 | -3.260 | 0.002 ** |
| NPRC (ENPRC) | -0.277 | 0.113 | -2.458 | 0.016 * |

**Note.** Model fit: quasibinomial GLM. Model including these predictors significantly improved fit compared to the intercept-only model ( $F_{3,101} = 9.96, p < 0.001$ ).

Across both centers, lower mean kinship predicted more severe aggression toward kin. NPRC also had a main effect: ENPRC matrilines directed proportionally less severe aggression toward kin than CNPRC matrilines. As expected, higher-ranking matrilines also used proportionally more severe aggression toward kin than lower-ranking ones.

#### Kinship Threshold Analysis

To evaluate whether kinship effects operated in a threshold rather than continuous manner, we re-fit the best-performing GLMs using binary predictors that categorized matrilines as either highly related or distantly related at cutoffs between 0.10 and 0.25. We applied this approach to models of grooming cohesion and severe aggression.

### Grooming

Threshold models consistently outperformed continuous kinship models. Across all variants, the strongest support was for a cutoff around 0.16 (Figure 3). For full grooming networks, the best-fit model for CNPRC groups identified a threshold of 0.16 (ΔAICc = –7), while ENPRC groups showed a slightly higher threshold (0.175) with negligible improvement over the continuous model (ΔAICc = –0.3). In the combined dataset, the 0.16 cutoff again provided the best fit (ΔAICc = –5.7). Analyses of density-matched networks yielded similar results, with threshold models favored in ENPRC (ΔAICc = –3.1), CNPRC (ΔAICc = –5.3), and combined datasets (ΔAICc = –8.1). These findings suggest that matrilineal cohesion is best predicted by a kinship threshold near 0.16.

**Figure 3.**
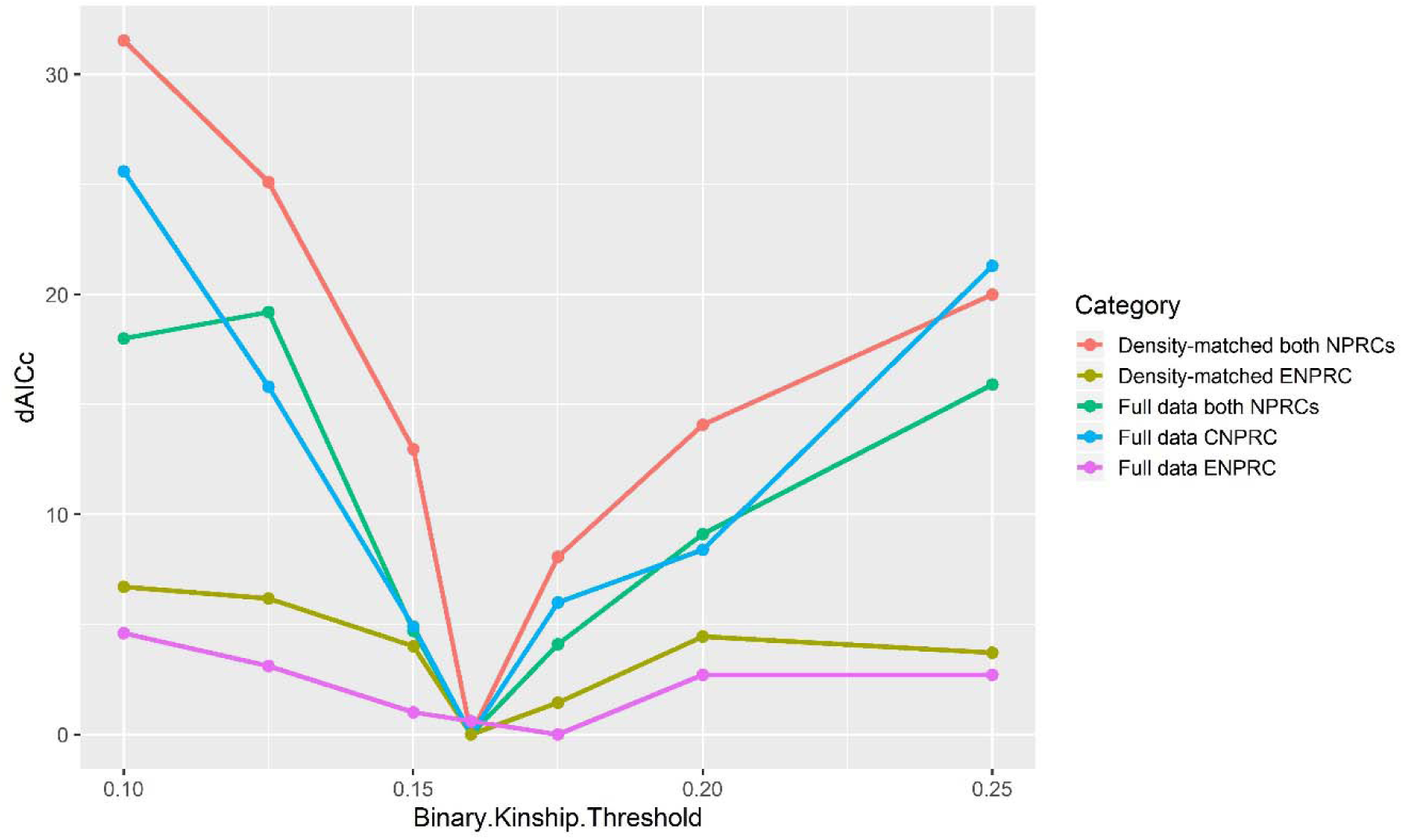
ΔAICc for GLMs with binary kinship thresholds (r = 0.10–0.25), relative to the best-fitting model. Model fit peaked near r = 0.16 across full and ENPRC density-matched datasets; CNPRC networks were not density-matched due to already low grooming density.

### Aggression

In contrast, threshold models did not improve fit for either aggression outcome. For the proportion of all aggression that was severe, the continuous kinship model (Residual Deviance = 308.97) provided the best fit, with all threshold models performing worse (Residual Deviance range: 347.74–381.76). Similarly, for the proportion of kin-directed severe aggression, threshold models did not outperform continuous kinship or reduced baseline models. These results indicate that, unlike grooming cohesion, severe aggression is not structured by a discrete kinship threshold.

## DISCUSSION

This study extends earlier work (Beisner et al., 2011b) by showing that matrilineal kinship remains a strong predictor of social cohesion in captive rhesus macaque groups, even across differences in housing conditions (group size) and management practices (presence/ absence of natal males). Matrilines with higher mean kinship consistently showed stronger grooming cohesion, with more female kin clustering together in the same grooming community. This pattern held true whether we accounted for variation in network density statistically or used density-matched networks. Because matrilineal alliances underpin female dominance hierarchies in rhesus macaques (Sade, 1969; Datta, 1986; Thierry, 2004), management practices for captive rhesus breeding groups should include routine monitoring of matriline size and mean kinship to identify potential genetic and social fragmentation that may warrant intervention.

To examine whether social cohesion weakens once families fall below a critical relatedness threshold, we tested a series of binary kinship models spanning a range of mean matriline kinship values. Across all model sets, the best-fitting threshold consistently fell between r = 0.16 and 0.175 — a range that aligns closely with previously identified dyadic thresholds for nepotistic bias (r = 0.25 and r = 0.125) (Kapsalis & Berman, 1996; Chapais et al., 1997; Silk, 2009). This consistent threshold suggests a predictable scaling effect, where deterioration of kin bias at the dyadic level corresponds to measurable declines in matriline-level cohesion.

Several factors may explain variation in matrilineal cohesion near this threshold. One is matriline composition: Chapais et al. (2001) found that in Japanese macaques, females intervened in conflicts involving *direct* kin (e.g., mother, grandmother) at relatedness levels as low as *r* = 0.125, whereas support for *collateral* kin (e.g., sisters, aunts) was typically limited to *r* ≥ 0.25. This suggests that matrilines with more direct kin may maintain stronger cohesion than those composed primarily of collateral relatives, even if average relatedness is similar.

These findings also align with Sherman’s ‘limits of nepotism’ hypothesis (Sherman, 1980, 1981), which offers an evolutionary explanation for reduced kinship among more distantly related individuals. Kin categories rarely encountered over evolutionary time—such as very distant or multigenerational relatives—may fall outside the bounds of evolved kin recognition mechanisms. In captive colonies, where large multigenerational matrilines can persist for many years, individuals may encounter kin types that would be rare or absent in the wild. This could explain why cohesion tends to deteriorate when average relatedness drops below *r* ≈ 0.16, as such kin may not be behaviorally distinguishable from non-kin.

At the same time, kinship is not the sole determinant of social bond strength. Affiliative preferences can also be shaped by shared rearing experiences, age proximity, or individual sociability. For instance, females often form particularly strong bonds with age-mates, regardless of kinship (Ehardt & Bernstein, 1987; Widdig et al., 2001; Silk et al., 2010; Liao et al., 2018). In some cases, highly cohesive relationships may form between more distantly related individuals due to unique social experiences—such as alloparental care or the absence of maternal support early in life. These patterns highlight the complexity of social dynamics in rhesus macaques: while kinship thresholds can identify matrilines at elevated risk of fragmentation, they do not determine which relationships will ultimately weaken or persist.

Based on time-constraint models and prior evidence from free-ranging and captive primates (Dunbar, 1992; Lehmann, Korstjens & Dunbar, 2007; Kaburu et al., 2019), we predicted that matrilines in larger groups would show lower grooming cohesion due to the cognitive and temporal costs of maintaining many social ties. Instead, we found a modest positive association between group size and matrilineal grooming cohesion, but only in the full network models—not in the density-matched analysis. This discrepancy suggests that the group size effect in full networks may reflect structural features, such as network density, that were more tightly controlled in the density-matched analysis. Interpretation is further complicated by uneven group sizes across centers: at CNPRC, groups ranged from 108 to nearly 200 animals, whereas most ENPRC groups had fewer than 100. Future studies should more explicitly disentangle these overlapping factors to clarify whether and how group size influences family cohesion.

While these findings underscore the importance of affiliative ties and kinship thresholds for predicting risk of fragmentation, the consequences of social breakdown are often more conspicuous in patterns of conflict. In particular, the use of severe aggression offers a revealing window into the erosion of cohesion within matrilines.

Whereas kinship thresholds shaped affiliative cohesion, aggression followed a different pattern: we found no evidence of a comparable threshold-like relationship. Instead, aggressive behavior scaled more continuously with kinship, suggesting that conflict dynamics are not organized around kinship thresholds in the same way as affiliation. As predicted, matrilines with lower mean kinship showed higher proportions of severe aggression overall, consistent with earlier findings (Beisner et al., 2011a). However, the strength and nature of the effect varied by facility and by the presence of natal males. At ENPRC, matrilines with lower mean kinship initiated more severe aggression, even in the absence of natal males. In contrast, at CNPRC, the same pattern emerged only when natal males were included in the analysis (Beisner et al., 2011a), underscoring that males can influence conflict dynamics in fragmented matrilines.

These cross-center differences emphasize the importance of natal male management. While their presence in the group allows them to leverage kin alliances to advance their own social status, which may undermine broader group stability (Beisner et al., 2011b), their removal can also destabilize group dynamics (Pritchard et al., 2024). Removing all natal males before they mature risks skewing sex ratios and reducing the pool of adult males available to police female conflicts (Beisner et al., 2011b). Retaining natal males may also provide long-term benefits by facilitating future breeding introductions (Rox et al., 2019). Thus, management strategies must balance the risks posed by natal males against the critical roles that adult males—particularly those unrelated to dominant matrilines—play in maintaining stability. At the same time, our results highlight a limitation: because our networks were defined by female kinship, we could not fully capture natal males’contributions to cohesion. Natal males forming affiliative bonds with mothers or sisters may help reinforce family ties, a possibility for future work. Nonetheless, matrilines with low mean kinship remain vulnerable to internal conflict regardless of natal male presence, underscoring the need for context-sensitive management.

A consistent pattern also emerged in the use of severe aggression among female relatives. Across both centers, matrilines with lower mean kinship directed proportionally more severe aggression toward kin. This finding suggests not only a breakdown in affiliative behavior and family cohesion, but also a shift in how females regulate their dominance relationships: in genetically fragmented families, females may rely less on milder forms of aggression and more on intense aggression to assert or defend their rank.

Kin frequently engage in low-level agonistic interactions, often due to spatial proximity and routine maintenance of dominance relationships (Kurland, 1977; Bernstein & Ehardt, 1985; Cheney & Seyfarth, 1999). Therefore, the elevated use of severe aggression among kin in low-relatedness matrilines seem to diverge from this norm, indicating that distant relatives may no longer be perceived as allies. In such contexts, females may increasingly view one another as competitors, particularly when affiliative bonds are insufficient to buffer rank-related tensions (Chapais, Savard & Gauthier, 2001; Silk, 2009; Beisner et al., 2011a).

These patterns are consistent with established models of kin recognition. Familiarity is the primary mechanism for kin recognition (Fredrickson & Sackett, 1984; Sackett & Fredrickson, 1987; Rendall, 2004). Early-life proximity to older siblings through maternal association facilitates the recognition of siblings as kin (Kapsalis & Berman, 1996; Berman & Kapsalis, 1999). However, as matrilines grow larger and more genetically fragmented, developmental familiarity may occur less consistently. Without reliable recognition cues, kin bias can break down, and females may treat distant relatives more like non-kin.

From a management perspective, this breakdown highlights the need for more sophisticated monitoring tools. Tracking matriline size, age structure, rearing histories, and mean kinship values could provide a more comprehensive assessment of social risk. The assumption that large, multigenerational matrilines will self-stabilize through kinship alone becomes increasingly tenuous in captive settings. This insight has implications not only for captive group management but also for understanding the limits of evolved social systems under novel demographic conditions.

Moreover, these dynamics have parallels in wild populations (Cheverud, Buettner-Janusch & Sade, 1978; Widdig et al., 2006). Kin-based subgrouping and eventual group fission have been documented in Japanese, bonnet, and toque macaques (Furuya, 1969; Koyama, 1970; Chepko-Sade & Sade, 1979; Dittus, 1988). The occurrence of similar fragmentation in natural settings underscores the robustness of these patterns and suggests that certain thresholds of kinship may universally shape group cohesion in this species.

### Study Limitations

Our findings should be interpreted in light of several limitations. First, group selection was not random: ENPRC groups were partly drawn from a male introduction study (only baseline data were used), groups lacking matrilineal structure were excluded, and some were chosen to broaden representation of larger groups at ENPRC to be more comparable to CNPRC groups. Second, our analyses focused on female family-level cohesion, which was appropriate for addressing matriline-level questions and facilitated cross-center comparison. However, this approach assessed natal male impacts only indirectly and did not consider alternative perspectives such as averaging across matriline members or analyzing all kin dyads. Finally, our study may not capture the full variation in social group structures that occur across colonies, given differences in kin composition and management practices. Thus, our results are most directly applicable to other captive breeding groups of similar composition, and extension to free-ranging populations should be cautious, as wild groups can resolve internal tensions through fission, a process unavailable to captive groups.

## Acknowledgments

This project was funded in part by the following NIH awards: P51 OD011132 to ENPRC, P51 OD011107 to CNPRC, R24 RR024396 to B. McCowan, R24 OD020349 to M. Bloomsmith, and R24 OD030035 to K. Ethun. We thank the animal care, colony management, and veterinary staff at the California and Emory NPRCs for their dedication to the management and welfare of the animals.

## Author Contributions

Brianne Beisner: Conceptualization (equal), Methodology (lead), Formal analysis (lead), Data curation (supporting), Visualization (lead), Writing – original draft (lead), Writing – review & editing (equal).

Brenda McCowan: Conceptualization (equal), Funding acquisition (lead), Data curation (equal),

Project administration (lead), Resources (lead), Supervision (lead), Writing – review & editing (equal).

Mollie Bloomsmith: Conceptualization (supporting), Funding acquisition (lead), Project administration (lead), Resources (lead), Supervision (lead), Writing – review & editing (equal).

Lauren Lacefield: Investigation (lead, ENPRC study), Data curation (lead, ENPRC study), Writing – original draft (supporting), Writing – review & editing (equal).

Kelly Ethun: Conceptualization (supporting), Funding acquisition (lead), Data curation (lead), Project administration (lead), Supervision (lead), Writing – review & editing (equal).

## Data Availability Statement

The data that support the findings of this study are available from the corresponding author upon request.

## Abbreviation

CNPRC: California National Primate Research Center
ENPRC: Emory National Primate Research Center

