## Supplementary Figures for "A cross-center comparison of the relationship between matriline fragmentation, grooming cohesion, and agonistic behavior in captive rhesus macaque (*Macaca mulatta*) social groups"

**Supplemental Information** – Beisner et al. Cross-Center Matriline Fragmentation

To evaluate how network density affects our measures of community structure, we conducted subsampling analyses for each ENPRC grooming network. For each group, we randomly reduced the data volume in 10% increments (from 90% to 10% of the original data, with 100 iterations per increment) and recalculated network modularity and the number of grooming communities. The resulting boxplots (Figures S1–S20) illustrate how these metrics vary as data density decreases. These analyses confirm that our density-matched networks reliably represent the underlying community structure of the full networks while allowing direct comparison with lower-density CNPRC networks.

**Figure S1.** Relationship between grooming network density and modularity for each NPRC. Lines show fitted trends from geom_smooth in *ggplot2*. Higher density is associated with lower modularity overall, motivating statistical control and density-matching

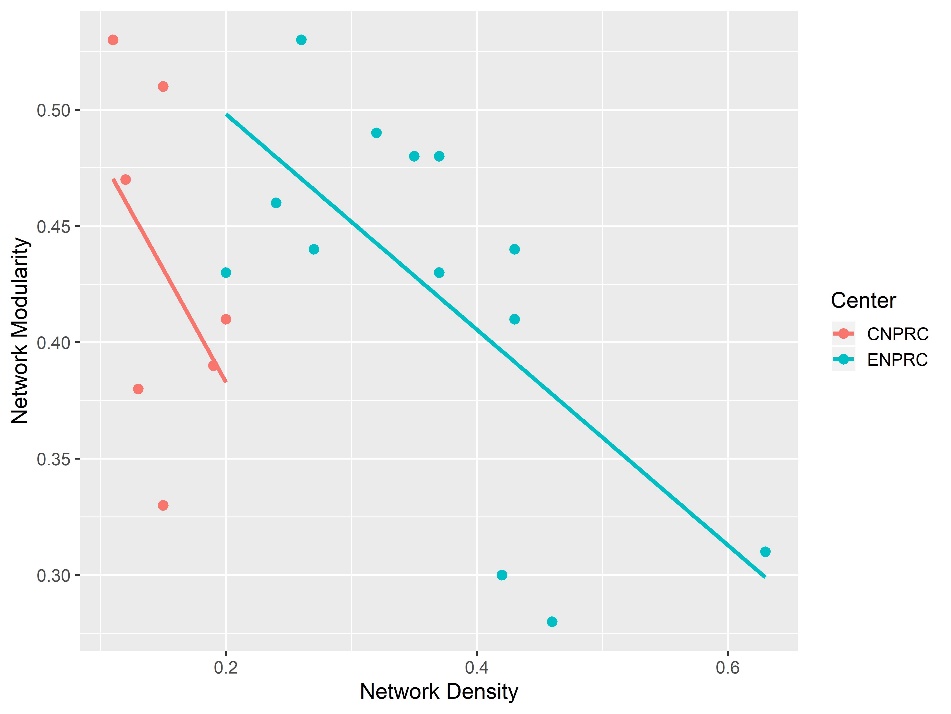

**Figure S2**. Change in Modularity by Subsampling Level for group II (study #2, phase 1)

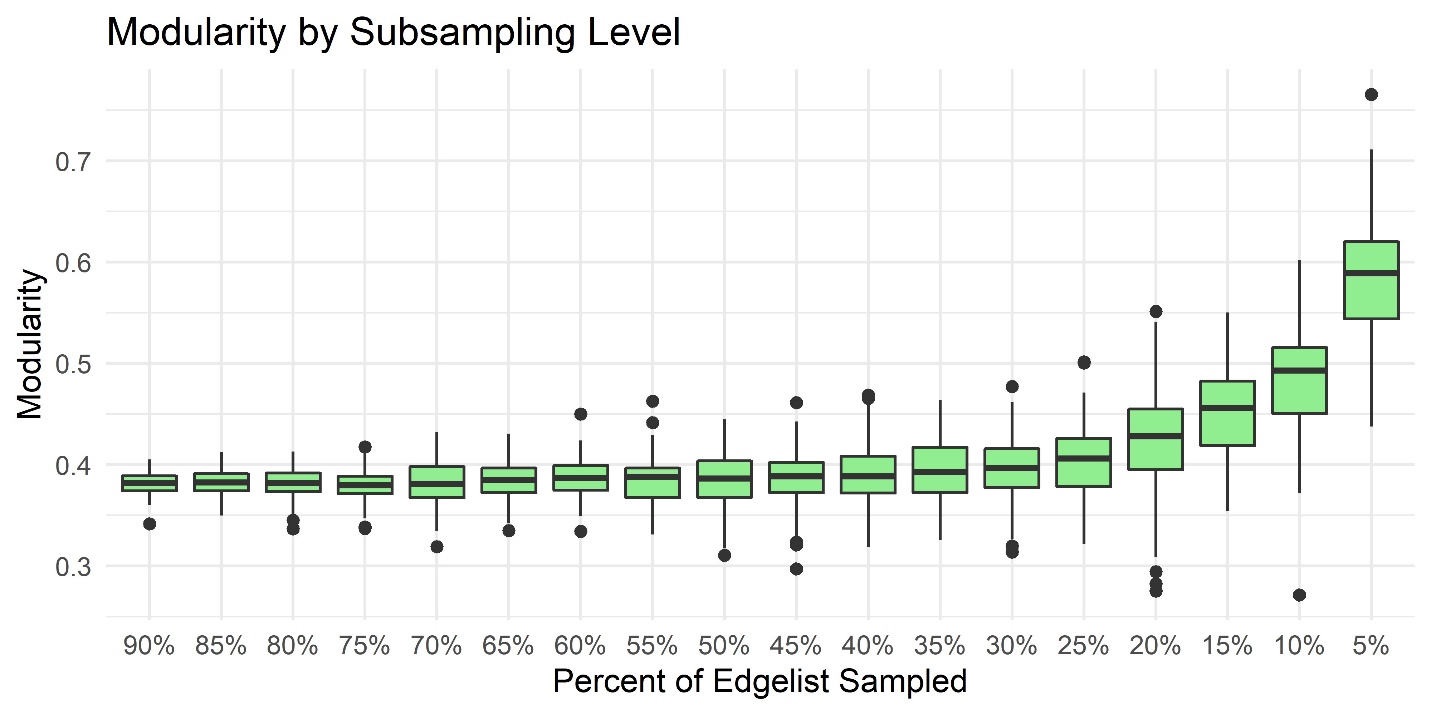

**Figure S3**. Change in Modularity by Subsampling Level for group VII (study #2, phase 3)

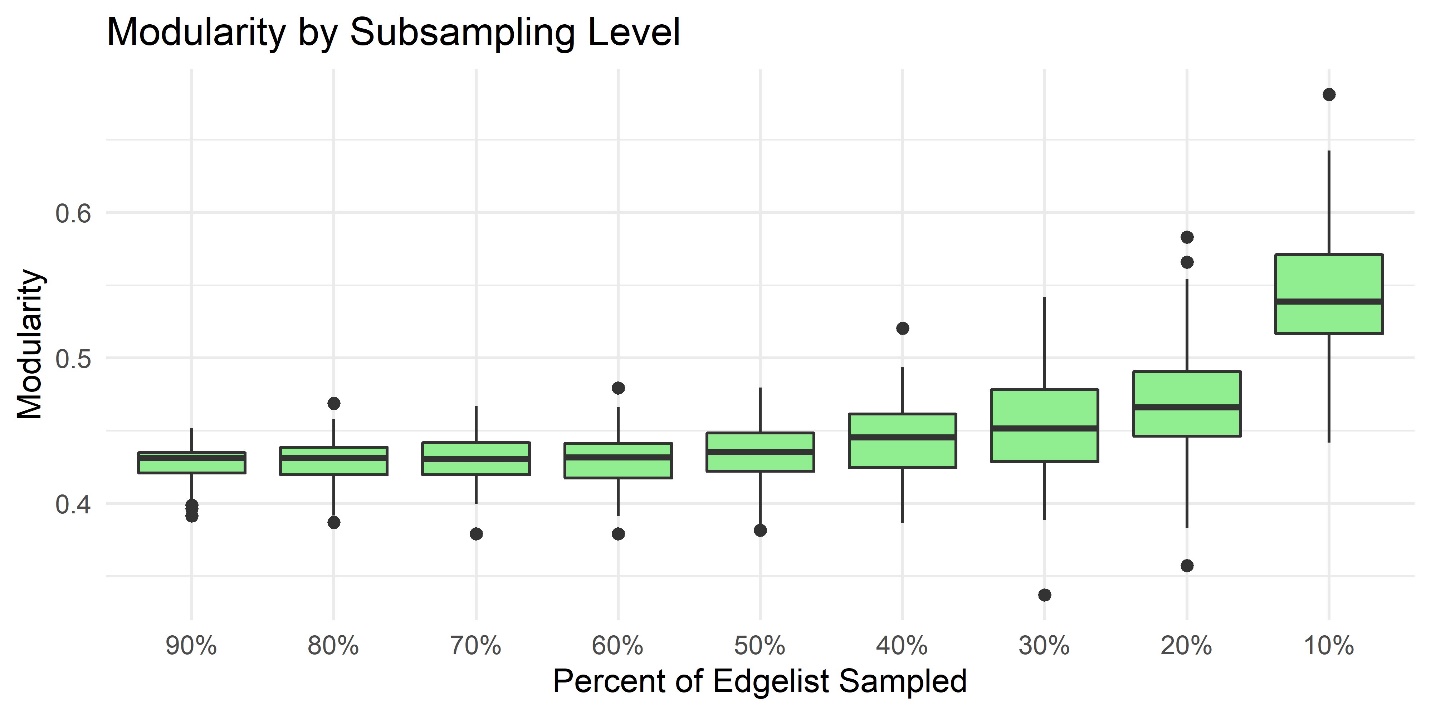

**Figure S4**. Change in Modularity by Subsampling Level for group V (study #2, phase 2)

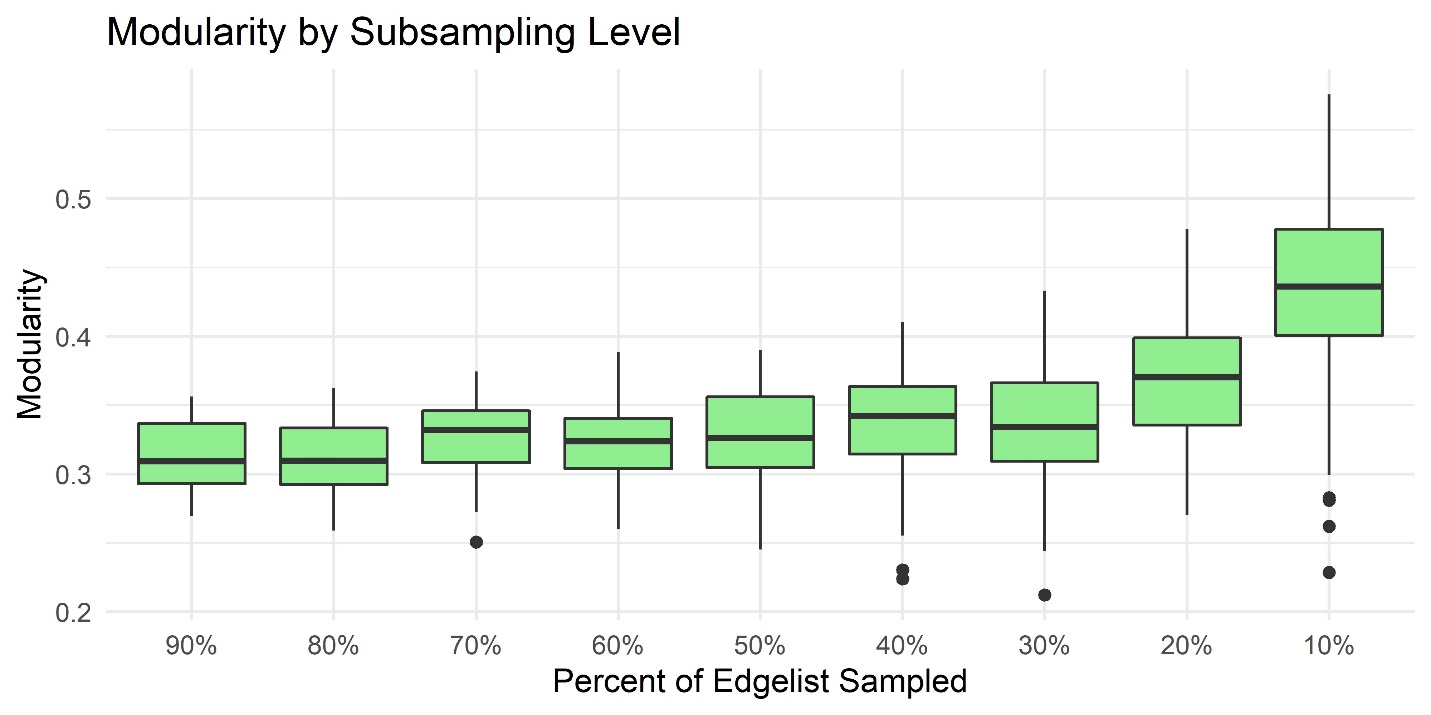

**Figure S5**. Change in Modularity by Subsampling Level for group 1 (study #2, phase 1)

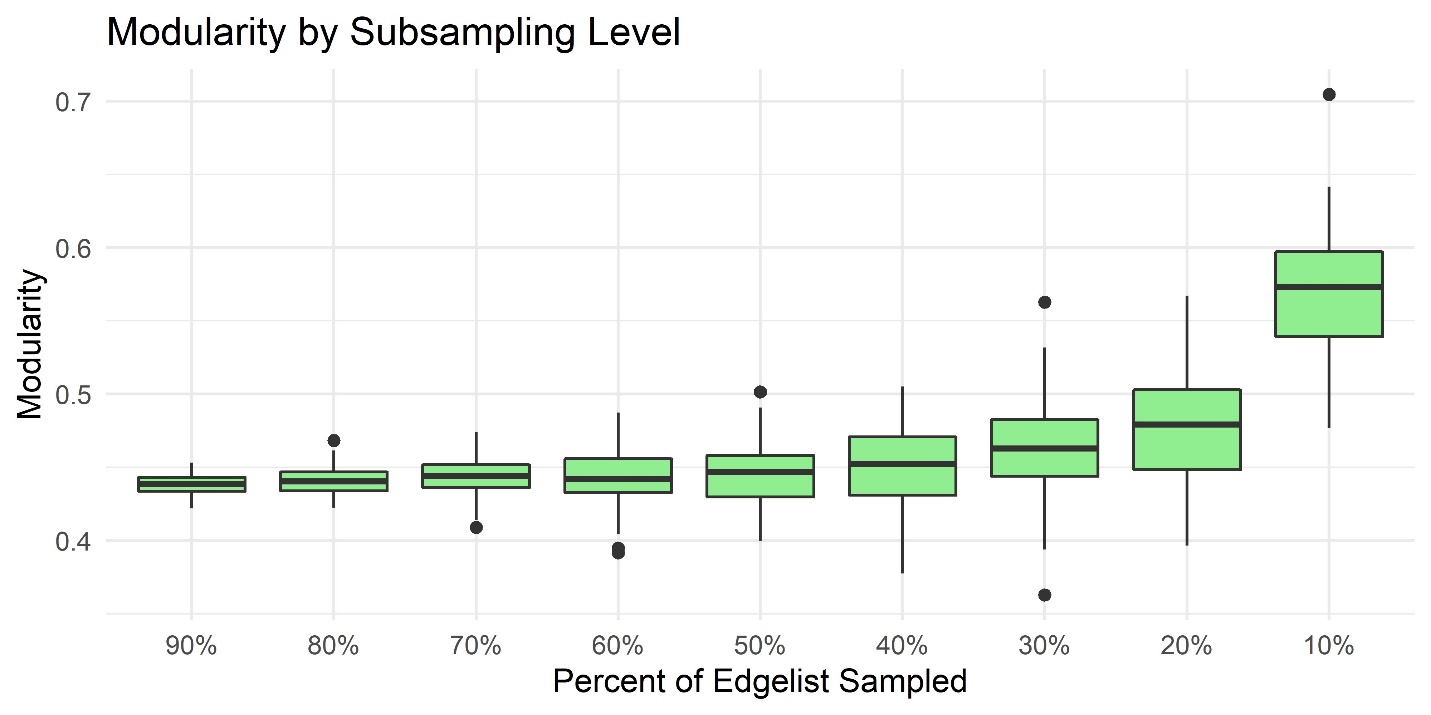

**Figure S6.** Change in Modularity by Subsampling Level for group VI (study #2, phase 2)

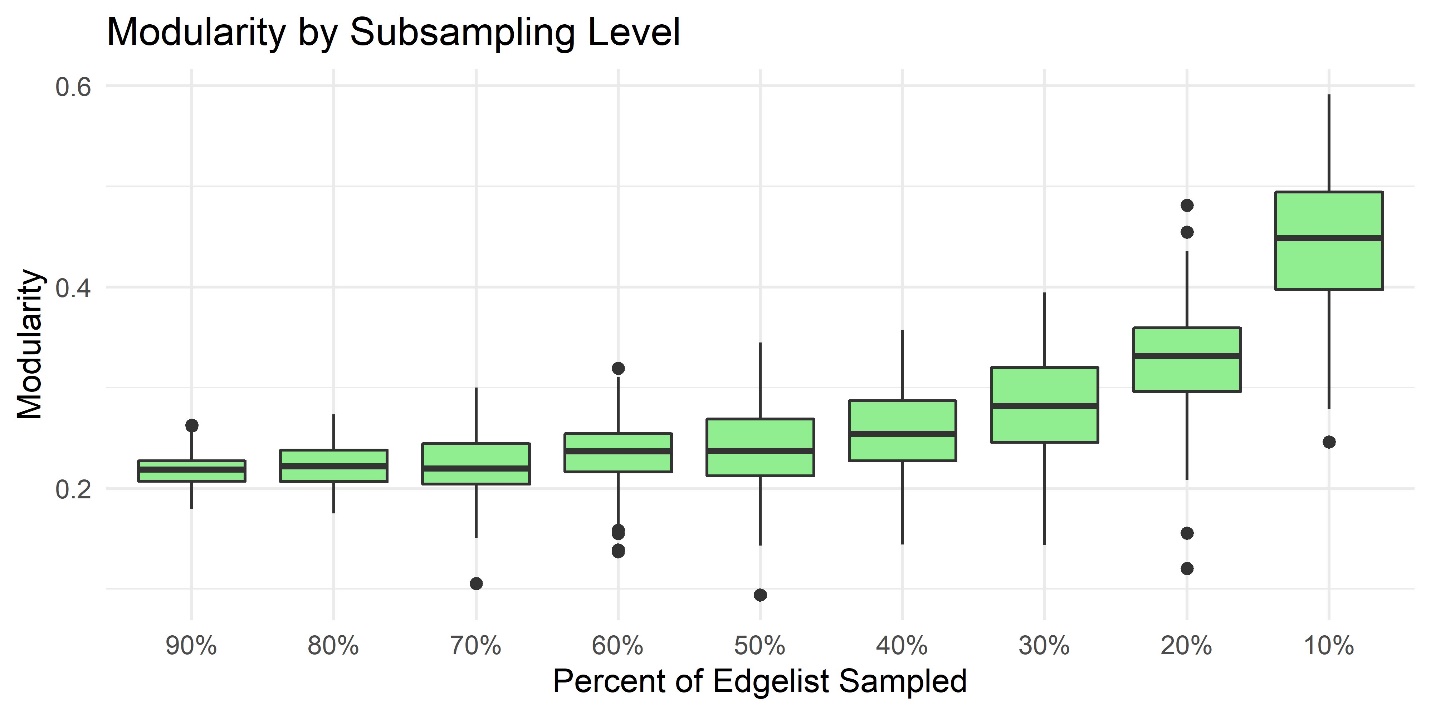

**Figure S7**. Change in Modularity by Subsampling Level for group II (study #2, phase 3)

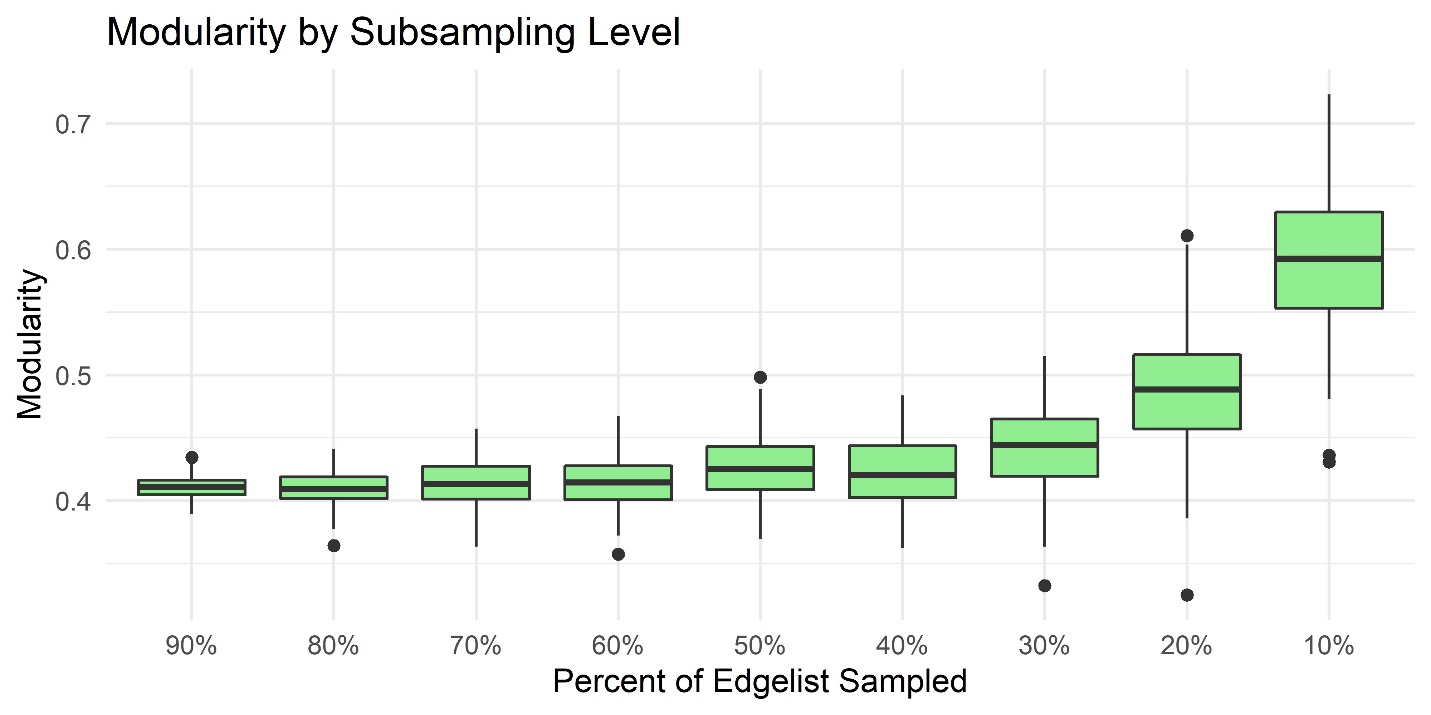

**Figure S8**. Change in Modularity by Subsampling Level for group IX (study #3)

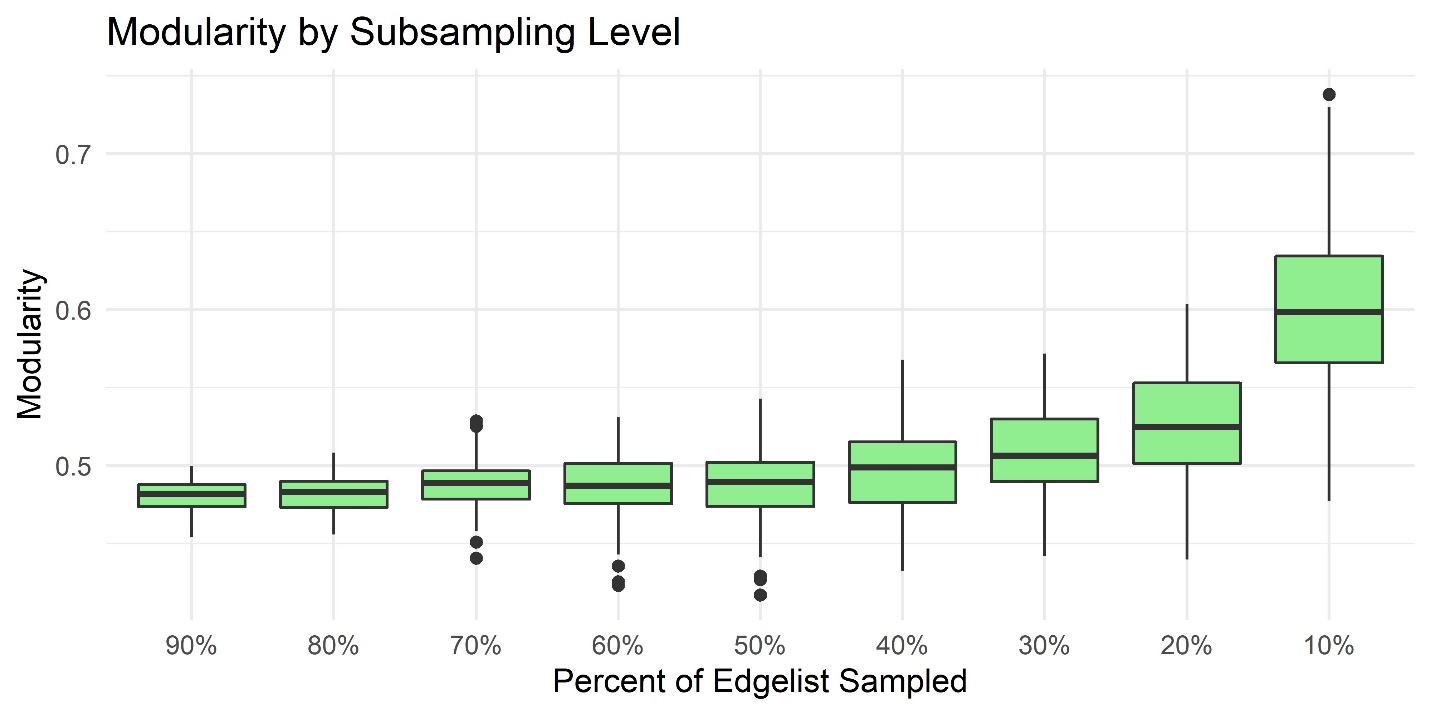

**Figure S9.** Change in Modularity by Subsampling Level for group VIII (study #3)

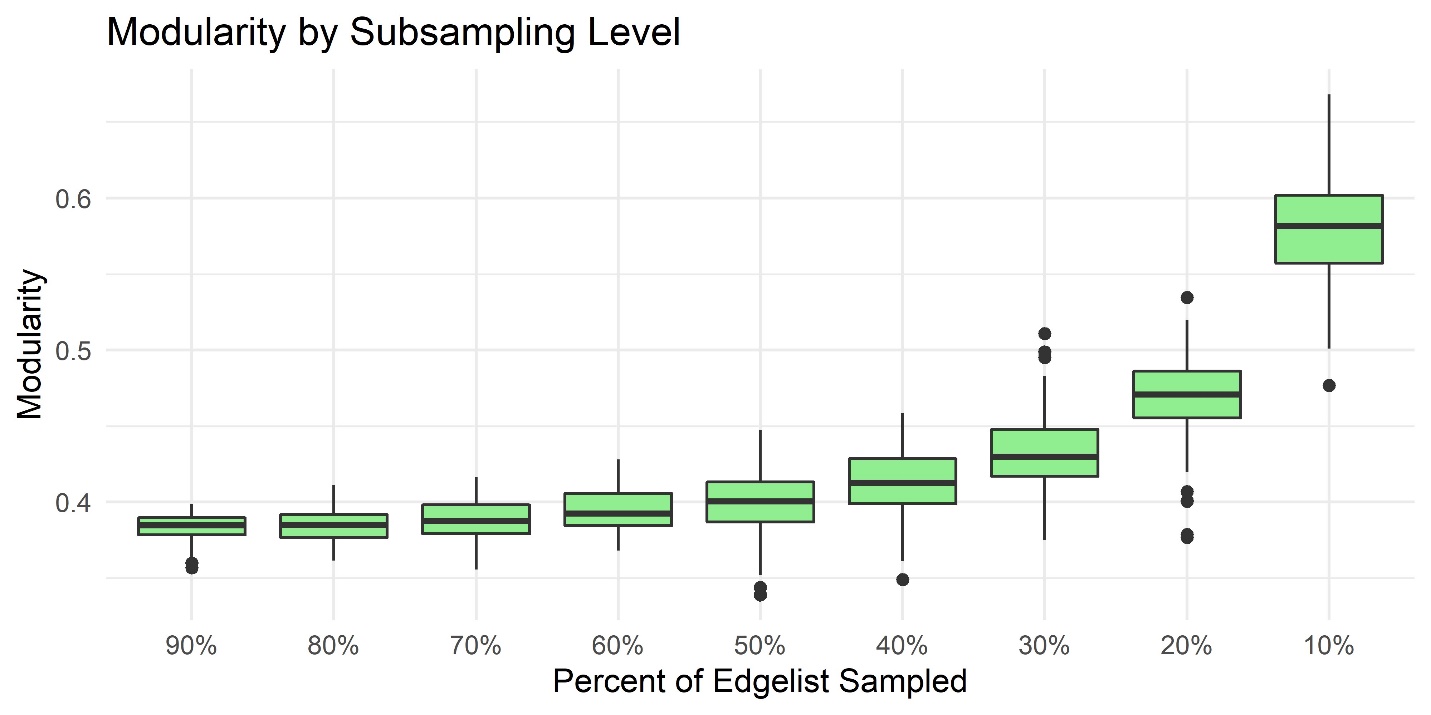

**Figure S10**. Change in Modularity by Subsampling Level for group X (study #3)
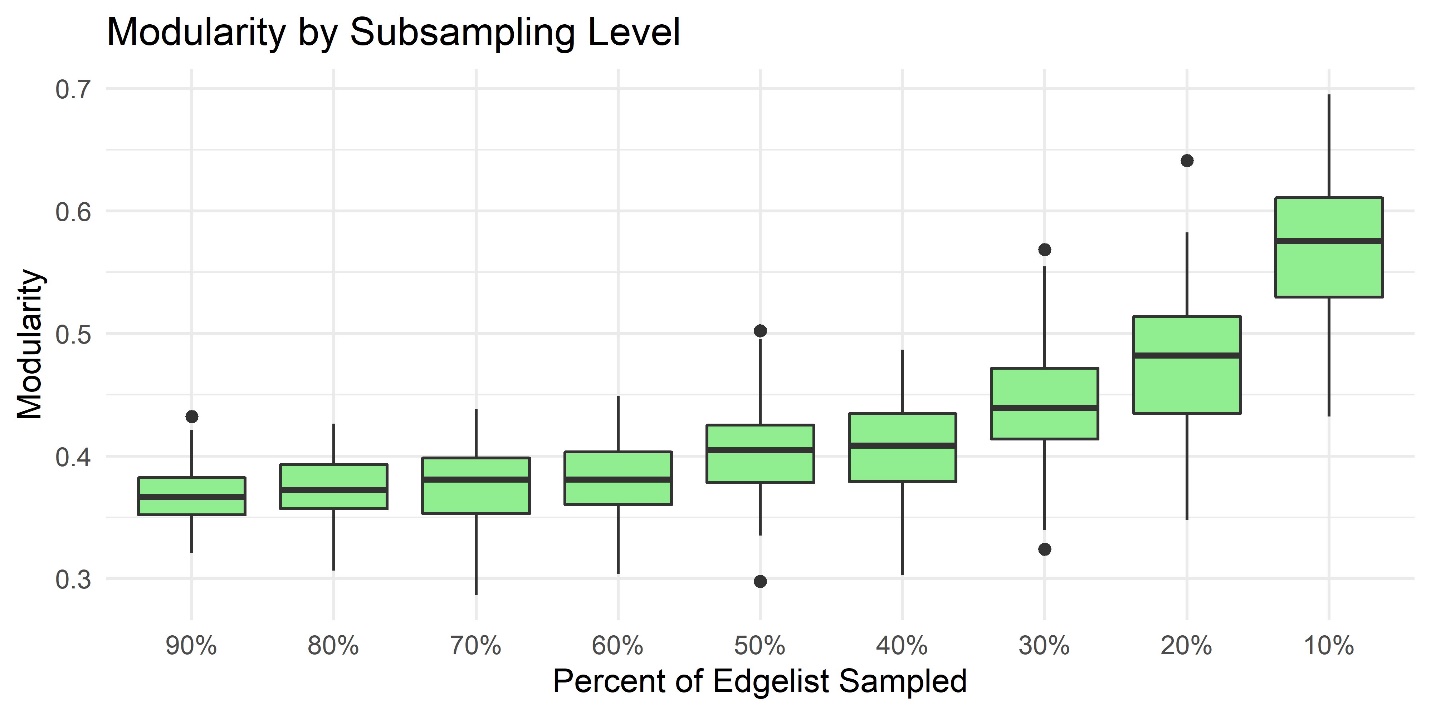

Boxplots of the number of groom network communities as edgelist data are subsampled from 90% to 10% for all ENPRC study groups.

**Figure S11.** Change in Number of Grooming Communities by Subsampling Level for group II (study #2, phase 1)

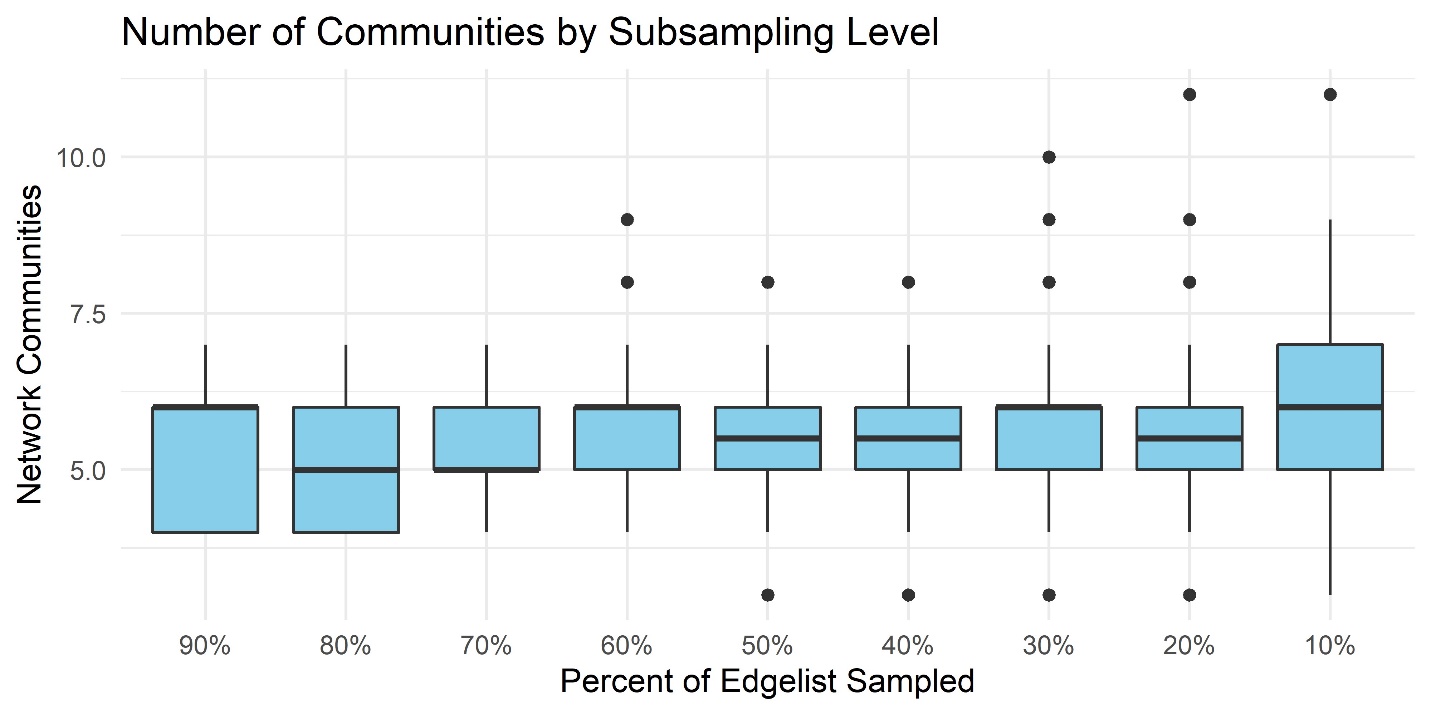

**Figure S12.** Change in Number of Grooming Communities by Subsampling Level for group VII (study #2, phase 3)

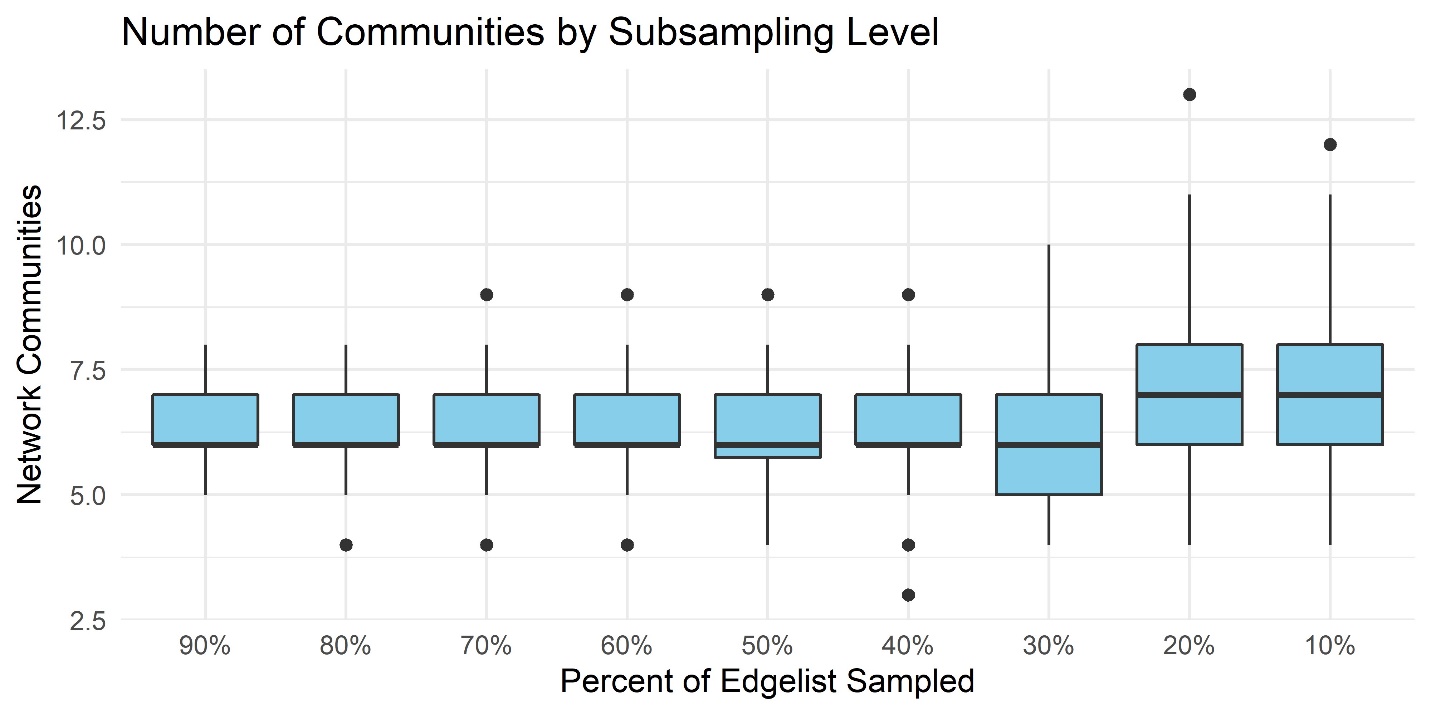

**Figure S13.** Change in Number of Grooming Communities by Subsampling Level for group V (study #2, phase 2)

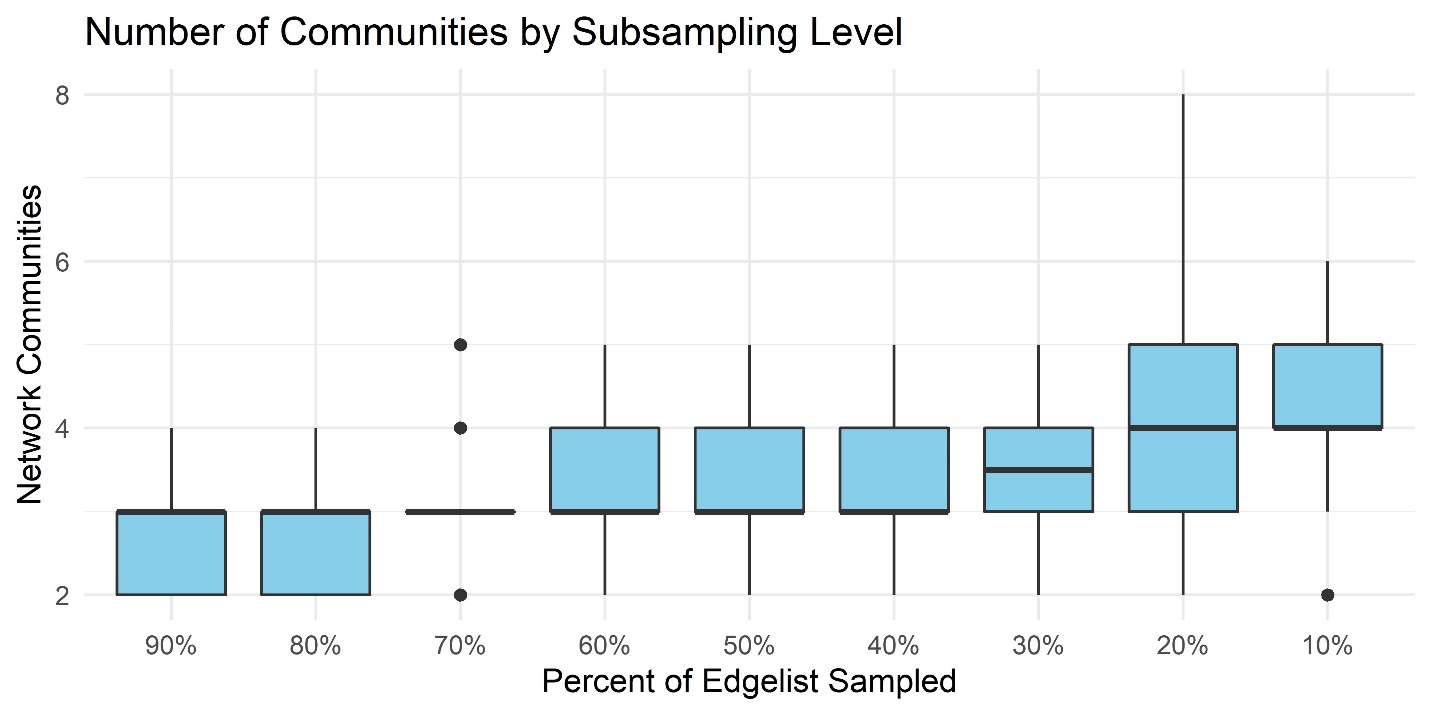

**Figure S14**. Change in Number of Grooming Communities by Subsampling Level for group I (study #2, phase 1)

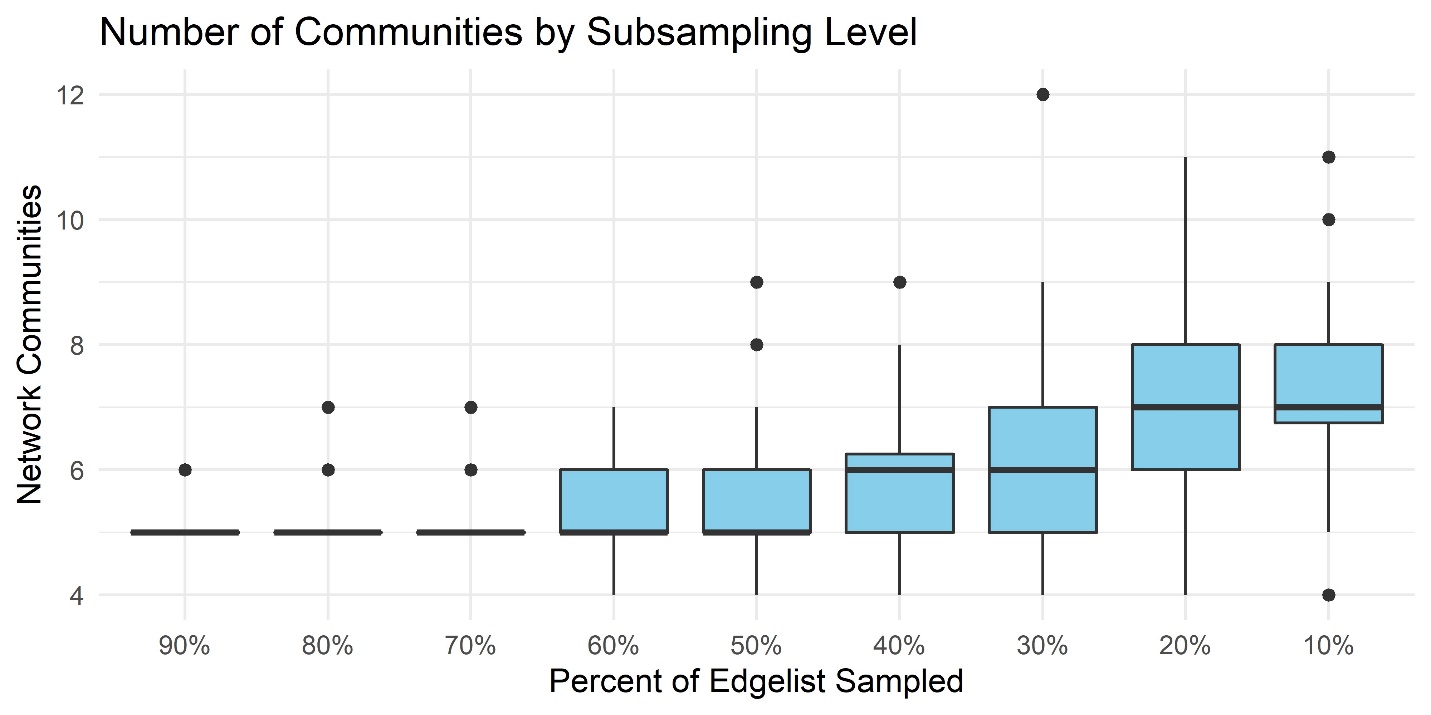

**Figure S15.** Change in Number of Grooming Communities by Subsampling Level for group II (study #2, phase 3)

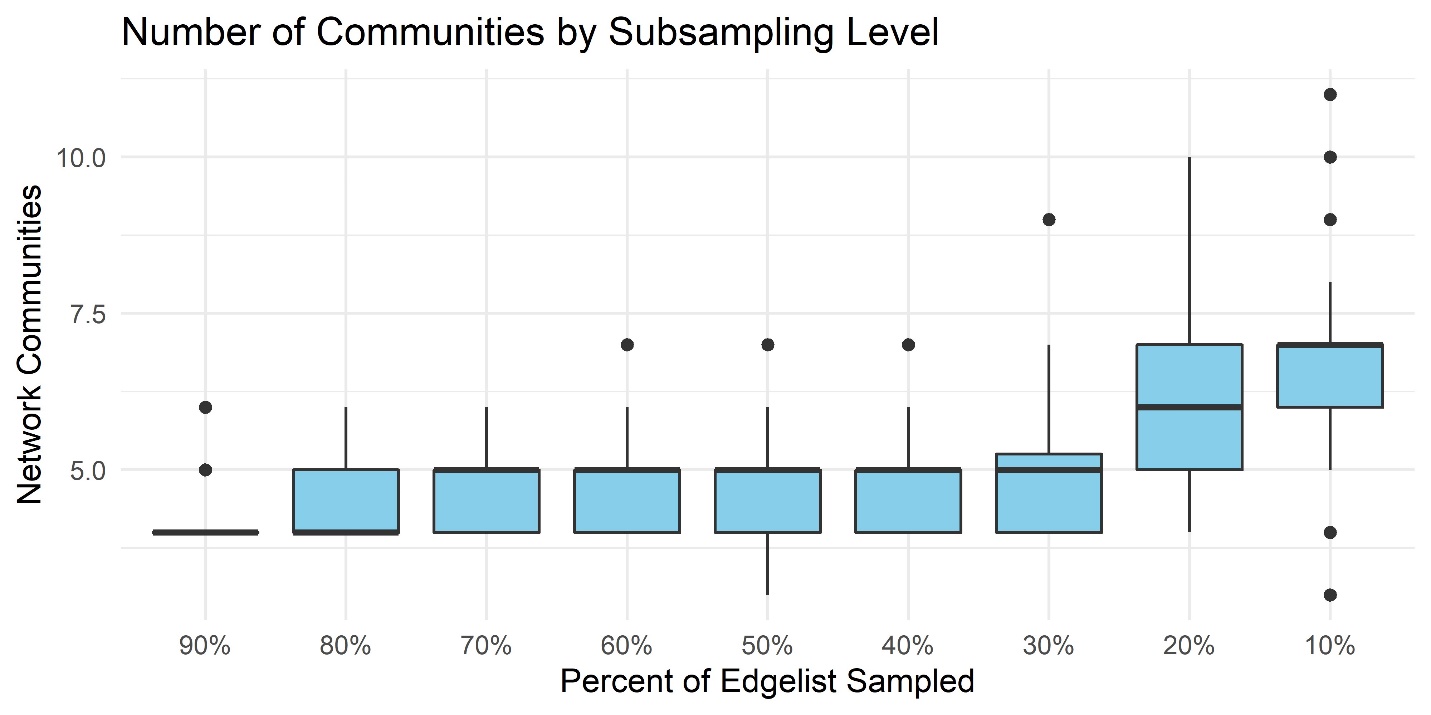

**Figure S16.** Change in Number of Grooming Communities by Subsampling Level for group VI (study #2, phase 2)

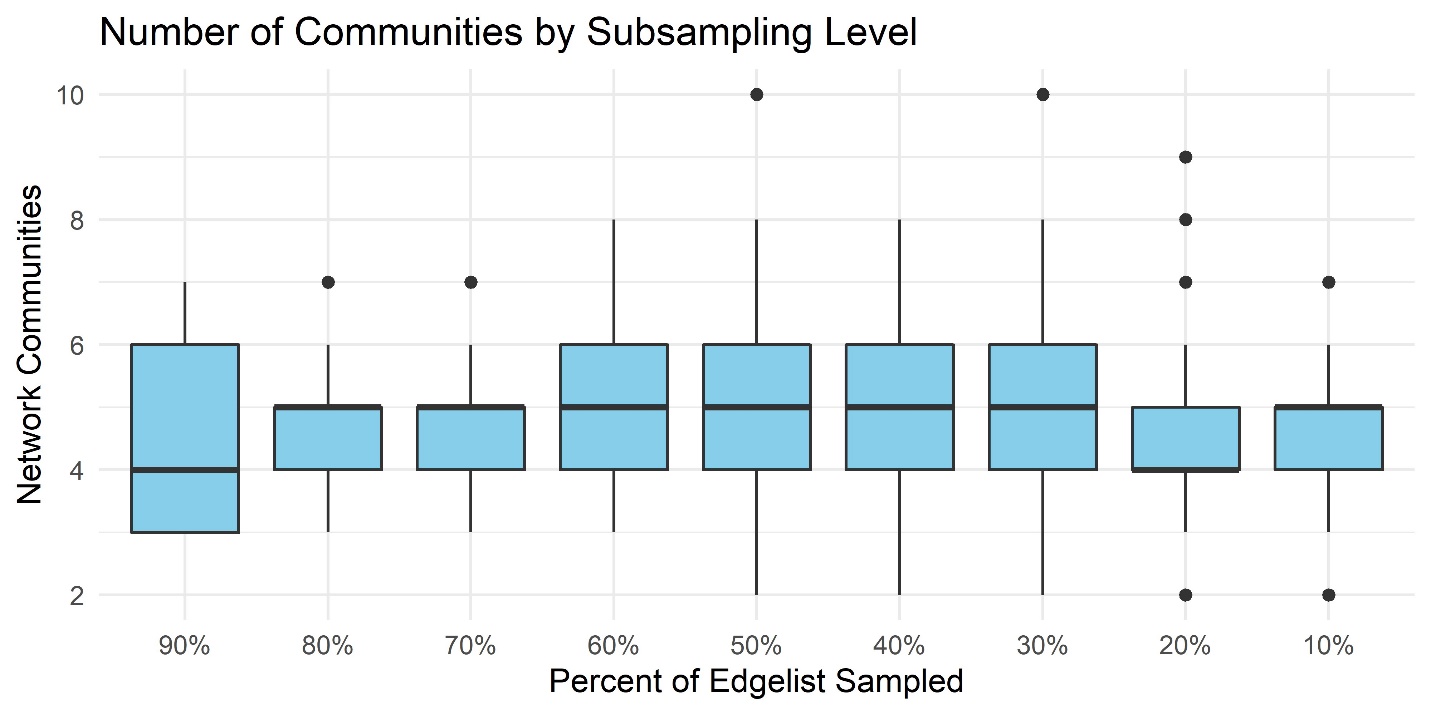

**Figure S17.** Change in Number of Grooming Communities by Subsampling Level for group II (study #3)

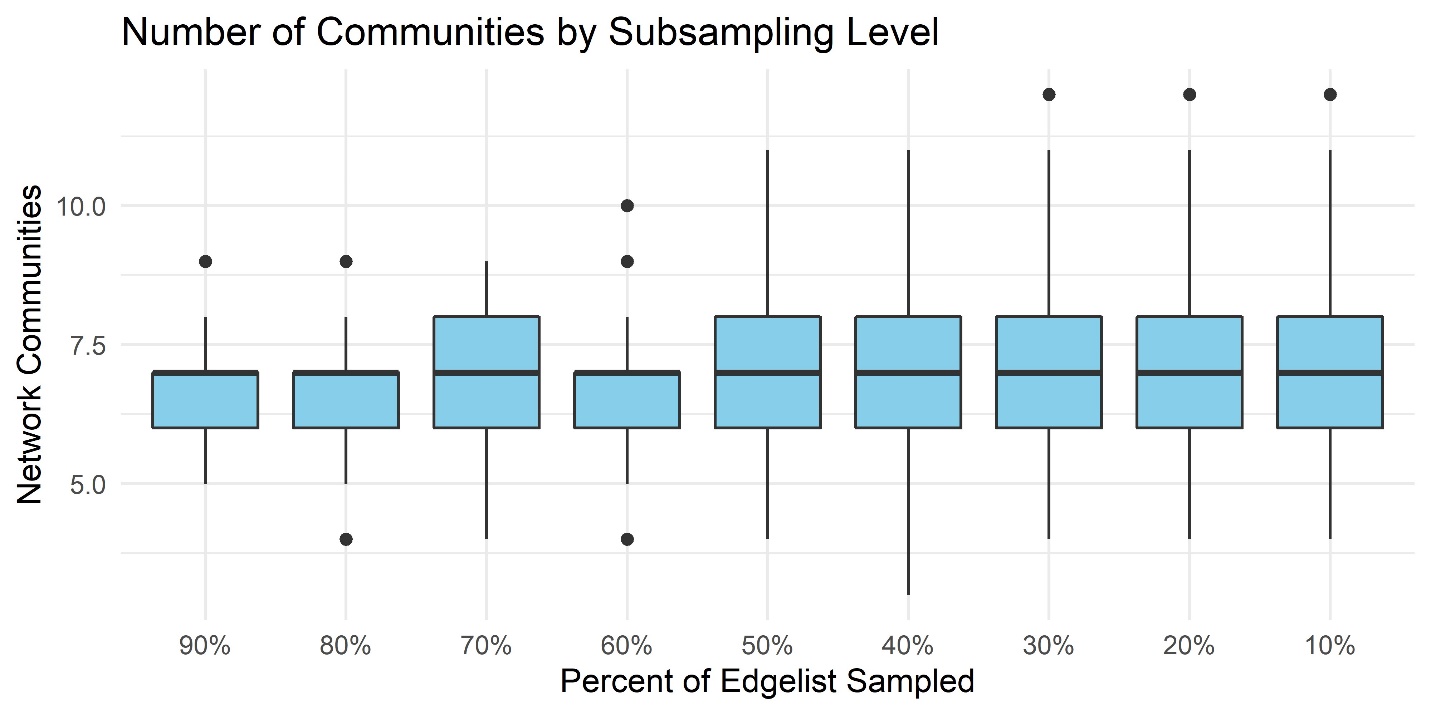

**Figure S18**. Change in Number of Grooming Communities by Subsampling Level for group IX (study #3)

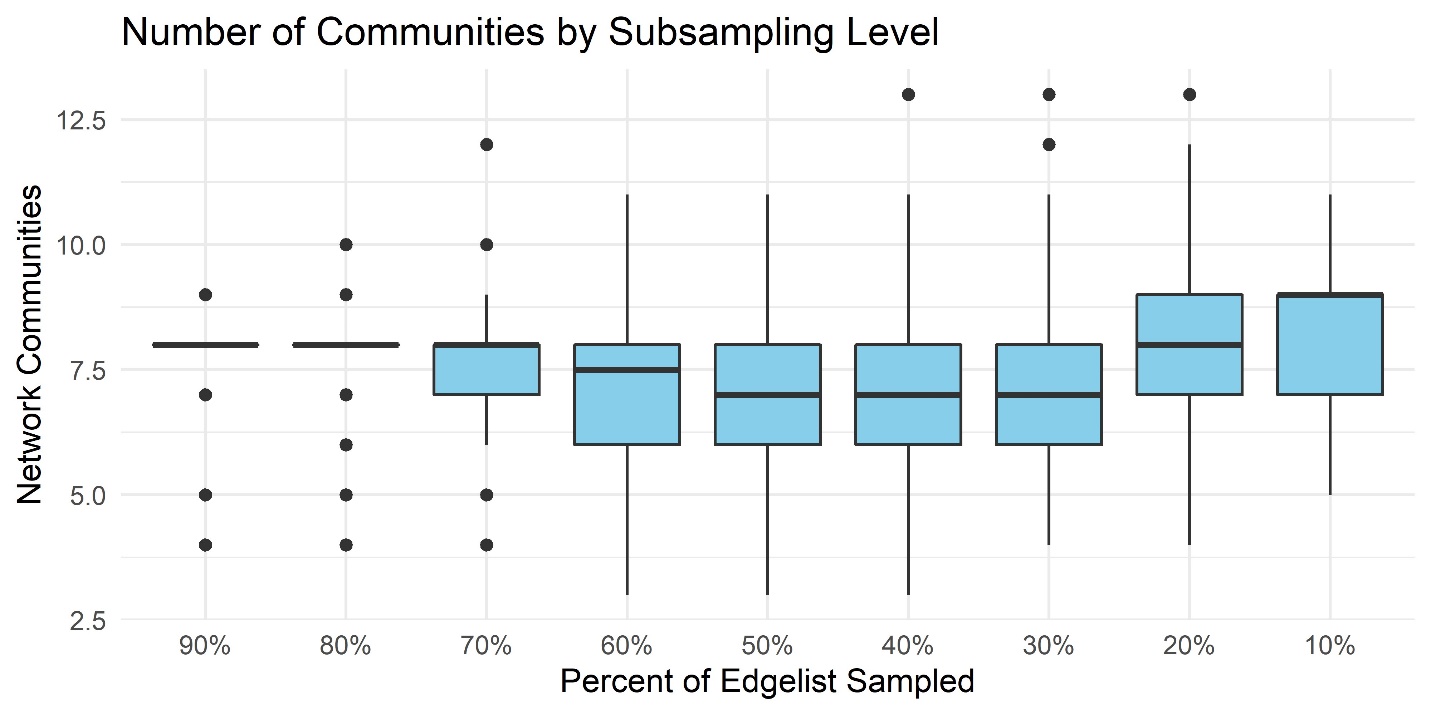

**Figure S19**. Change in Number of Grooming Communities by Subsampling Level for group VIII (study #3)

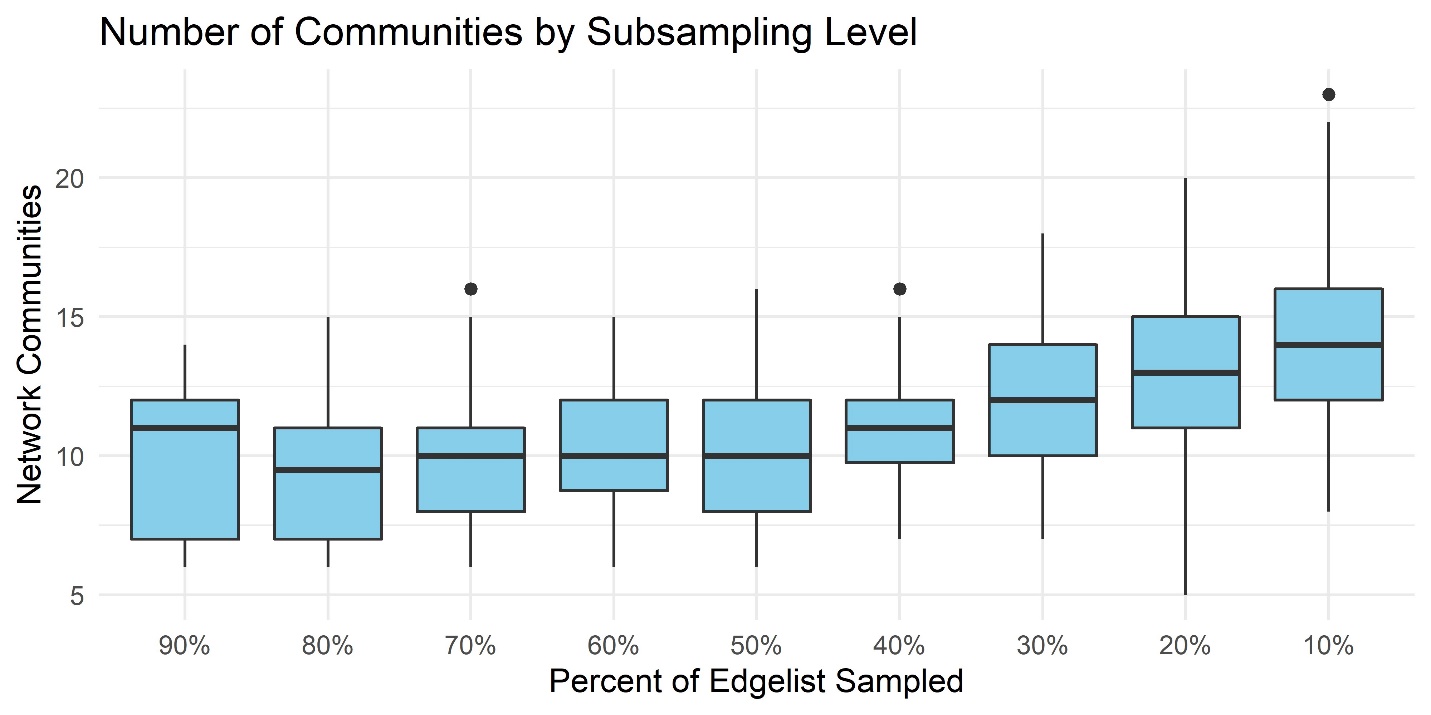

**Figure S20**. Change in Number of Grooming Communities by Subsampling Level for group X (study #3

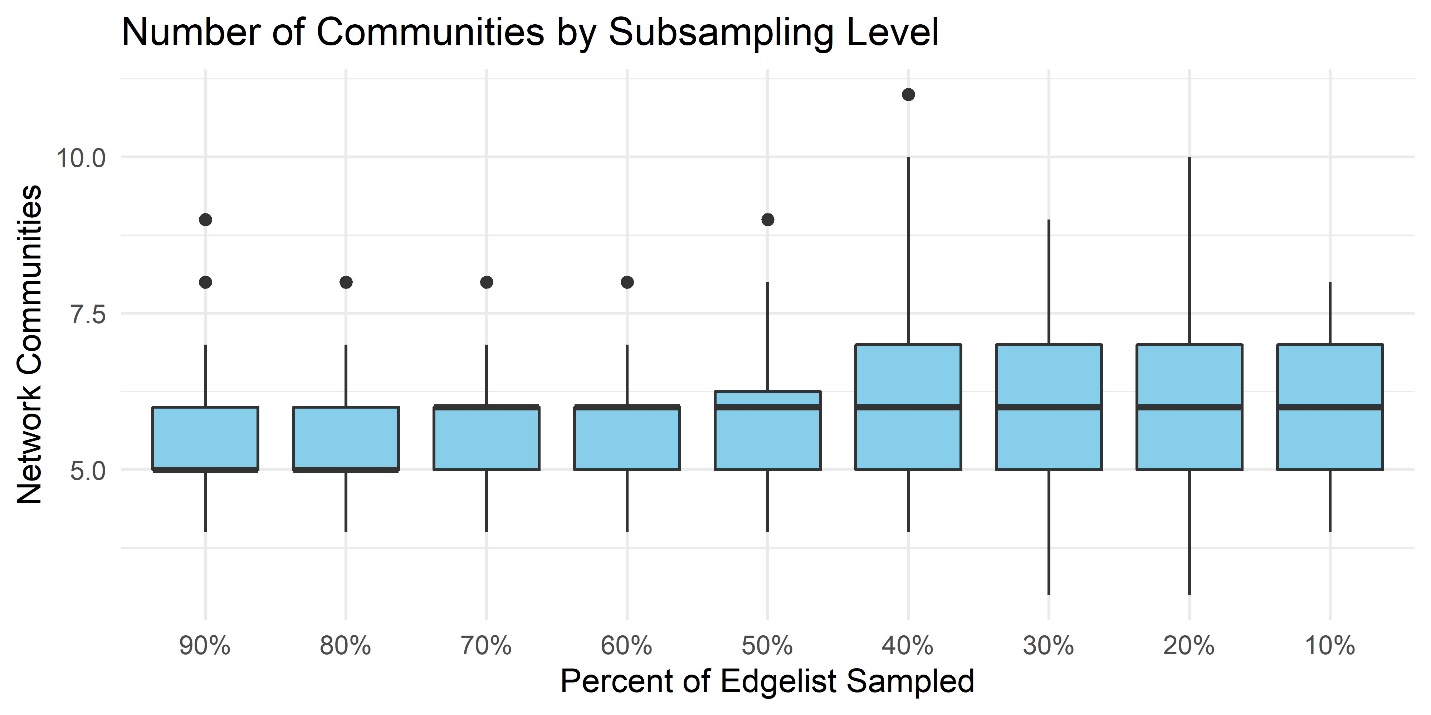
